# The *GPAT4*/*6*/*8* clade functions in Arabidopsis root suberization non-redundantly with the *GPAT5/7* clade required for suberin lamellae

**DOI:** 10.1101/2024.03.13.584743

**Authors:** Kay Gully, Alice Berhin, Damien De Bellis, Cornelia Herrfurth, Ivo Feussner, Christiane Nawrath

## Abstract

**Summary:** Lipid polymers such as cutin and suberin strengthen the diffusion barrier properties of the cell wall in specific cell types and are essential for water relations, mineral nutrition, and stress protection in plants. Land plant–specific glycerol-3-phosphate acyltransferases (GPATs) of different clades are central players in cutin and suberin monomer biosynthesis. Here, we show that the *GPAT4*/*6*/*8* clade in *Arabidopsis thaliana*, which is known to mediate cutin formation, is also required for developmentally regulated root suberization, in addition to the established roles of *GPAT5/7* in suberization. The *GPAT5*/*7* clade is mainly required for abscisic acid–regulated suberization. In addition, the *GPAT5*/*7* clade is crucial for the formation of the typical lamellated suberin ultrastructure observed by transmission electron microscopy, as distinct amorphous globular polyester structures were deposited in the apoplast of the *gpat5 gpat7* double mutant, in contrast to the thinner but still lamellated suberin deposition in the *gpat4 gpat6 gpat8* triple mutant. The intrinsic phosphatase activity of GPAT4, GPAT6, and GPAT8, which leads to monoacylglycerol biosynthesis, may be important for suberin biosynthesis. GPAT5/7 lack an active phosphatase domain. Notably, *gpat5 gpat7* phenotypes were partially reverted by treatment with a phosphatase inhibitor or the expression of phosphatase-dead variants of *GPAT4*/*6*/*8.* Thus, GPATs that lack an active phosphatase domain, which are predicted to synthetize lysophosphatidic acids, might be crucial for the formation of the lamellated structure of suberin. GPATs with active and non-active phosphatase domains appear to have non-redundant functions and must cooperate to achieve the efficient biosynthesis of correctly structured suberin.

**Significance statement:** The establishment of proper lamellated suberin in roots plays essential roles in regulating mineral nutrition and water relations in plants. The basis for the macromolecular arrangement determining the ultrastructure and properties of suberin lamellae is unknown. Here, we report that both the *GPAT4*/*6*/*8* and *GPAT5*/*7* clades of glycerol-3-phosphate acyltransferases (GPATs) contribute to suberin formation in Arabidopsis roots. In addition, we reveal that the *GPAT5*/*7* clade is required for the formation of the lamellated suberin ultrastructure. Several lines of evidence suggest that the loss of phosphatase activity in GPATs might have played a crucial role in the formation of suberin lamellae during evolution.

## Introduction

A crucial event in the evolution of land plants was the development of polymers rich in aliphatic and/or aromatic compounds, such as cutin, suberin, and lignin. These hydrophobic polymers strengthen the polysaccharide-rich cell walls and/or improve their diffusion barrier properties, enabling plants to live on land under fluctuating environmental conditions (1, 2).

Suberin is deposited as a secondary cell wall structure in specific tissues during plant development. For example, suberin is deposited in the endodermis and exodermis of the root, as well as during periderm and seed coat formation when tissues must be sealed off from the outside environment (3, 4). In addition, suberin is laid down in response to certain stress conditions, such as wounding or abscisic acid (ABA)–regulated abiotic stress, even in tissues that typically do not suberize, such as the root cortex or fruit epidermis; the latter leads to rough skin on fruits, known as russeting (5).

Aliphatic and aromatic components can be released from suberized cell walls by ester-bound cleaving agents. The partial depolymerization of suberin has revealed a network of aliphatic compounds, glycerol, and aromatic components in suberin (4). The suberin polyester is tightly associated with a lignin-like polymer in which the aromatic monomers are linked by C-C and C-O-C bonds and mainly contains hydroxycinnamates (mostly ferulate), instead of the monolignols found in lignin (6). Due to the tight association of the suberin polyester with a lignin-like polymer, suberin can be defined as a complex polymer with a polyaliphatic and polyaromatic domain (7). Other researchers prefer to characterize suberin as a largely aliphatic polyester similar to cutin associated with a lignin-like polymer (2, 8). The latter definition of suberin was adopted in this article.

The suberized cell wall forms a typical lamellar ultrastructure of electron-lucent (light) and electron-opaque (dark) lamellae, which were hypothesized to consist of the aliphatic suberin polyester and a polyaromatic polymer, respectively (4, 9). Despite the importance of ferulate for endodermal barrier formation (10), mutant analyses suggested that ester-bound ferulate is not essential for the formation of the suberin lamella ultrastructure in Arabidopsis (*Arabidopsis thaliana*) and potato (*Solanum tuberosum*) (11–13). However, the knockout of *AtCYP86A1* or its ortholog in potato led to distortions and altered dimensions of the suberin lamellae, indicating that aliphatic suberin monomers contribute to suberin lamellae formation (12, 14).

Suberin shares many compositional similarities with cutin. Both polyesters are rich in oxygenated fatty acid derivatives, such as hydroxy and/or epoxy acids, diacids, and fatty alcohols, but they also contain unsubstituted fatty acids. In addition to monomers with a chain length of C16 and C18, suberin typically contains more C20–C26 monomers than cutin. Glycerol and hydroxycinnamic acids are typically more abundant in suberin, and ferulate is a suberin component that is uncommon in cutin (15, 16).

Suberin is formed by the same polyester biosynthetic pathway as cutin, but often by different members of the same protein families (2, 17, 18). The land plant–specific family of glycerol-3-phosphate acyltransferases (GPATs), which is subdivided into three clades, plays a central role in the biosynthesis of polyesters (19). Members of two closely related clades, GPAT4/GPAT6/GPAT8 and GPAT5/GPAT7, localize to the endoplasmic reticulum (ER), as shown experimentally for GPAT8 (20), while enzymes of the GPAT1/GPAT2/GPAT3 clade appear to localize to mitochondria (21, 22). All land plant–specific GPATs have *sn2* specificity for the transfer of the acyl chain to glycerol-3-phosphate (19). GPATs have a phosphatase domain, in addition to their acyltransferase domain. However, only GPAT4, GPAT6, and GPAT8 have a phosphatase domain that has retained its typical active center and synthesizes monoacylglycerols (MAGs) (19, 23). The active site was not maintained during the evolution of other GPATs, including GPAT5 and GPAT7, resulting in lysophosphatidic acids (LPAs) as products (23). The difference in GPAT enzymatic activity has been hypothesized to distinguish cutin and suberin biosynthesis (17, 24). Arabidopsis GPAT4 and GPAT8 act redundantly in cutin formation in leaves and stems (25), while Arabidopsis GPAT6 is required for cutin formation in flowers as well as polyester formation (sporopollenin) in pollen (26, 27). GPAT6 was recently found to contribute to suberin formation as well (28).

Despite the similarities between cutin and suberin biosynthesis, distinct molecular mechanisms underlie their different positions in the apoplast (e.g., suberin in the secondary cell wall and cutin in and on the surface of the primary cell wall), the relationship between the suberin polyester and a polyaromatic polymer, and the formation of the characteristic suberin lamellae, remain unclear.

Here, we report that *GPAT4, GPAT6,* and *GPAT8* contribute non-redundantly to root suberization in addition to *GPAT5* and *GPAT7.* Analysis of the clade-specific Arabidopsis *gpat5 gpat7* and *gpat4 gpat6 gpat8* knockout mutants revealed that the *GPAT5/7* clade is required for the formation of the typical suberin lamellae since in the *gpat5 gpat7* double mutant suberin was deposited as discrete globular, amorphous structures. Mutating the active sites of the phosphatase domain in GPAT4, GPAT6, and GPAT8 reduced the amount of lamellated suberin. Expressing the genes encoding phosphatase-dead versions of GPAT4, GPAT6, and GPAT8 in the *gpat5 gpat7* double mutant resulted in the formation of a continuous suberin layer including lamellae. This study reveals distinct roles of GPATs with and without phosphatase domain in suberin lamellae formation.

## Results

### Expression patterns of *GPAT5/7* and *GPAT4/6/8* clade *GPAT*s in the endodermis

The land plant–specific GPATs required for polyester biosynthesis form three different clades (Fig. 1*A*) (19, 29). To examine the cell type–specific expression of *GPAT1* to *GPAT8*, we constructed promoter*GPATx:nls-GFP-GUS* transcriptional reporters (with the *GPATx* promoter driving the expression of a fusion construct encoding green fluorescent protein fused to β-GLUCURONIDASE with a nuclear localization sequence) and introduced them into wild-type (WT) Arabidopsis plants. We evaluated *GPAT* expression patterns in the roots of 5-day-old seedlings based on GFP fluorescence. In addition to the known *GPAT5* expression in the root endodermis (30, 31), its close homolog *GPAT7,* which was hypothesized to act in suberin formation, also showed endodermal expression (Fig. 1*B*) (17, 19). *GPAT4*, *GPAT6*, and *GPAT8* were also expressed in the root endodermis (Fig. 1*B*), unlike the members of the *GPAT1*/*2*/*3* clade (*SI Appendix*, Fig. S1*A*).

**Fig. 1.**
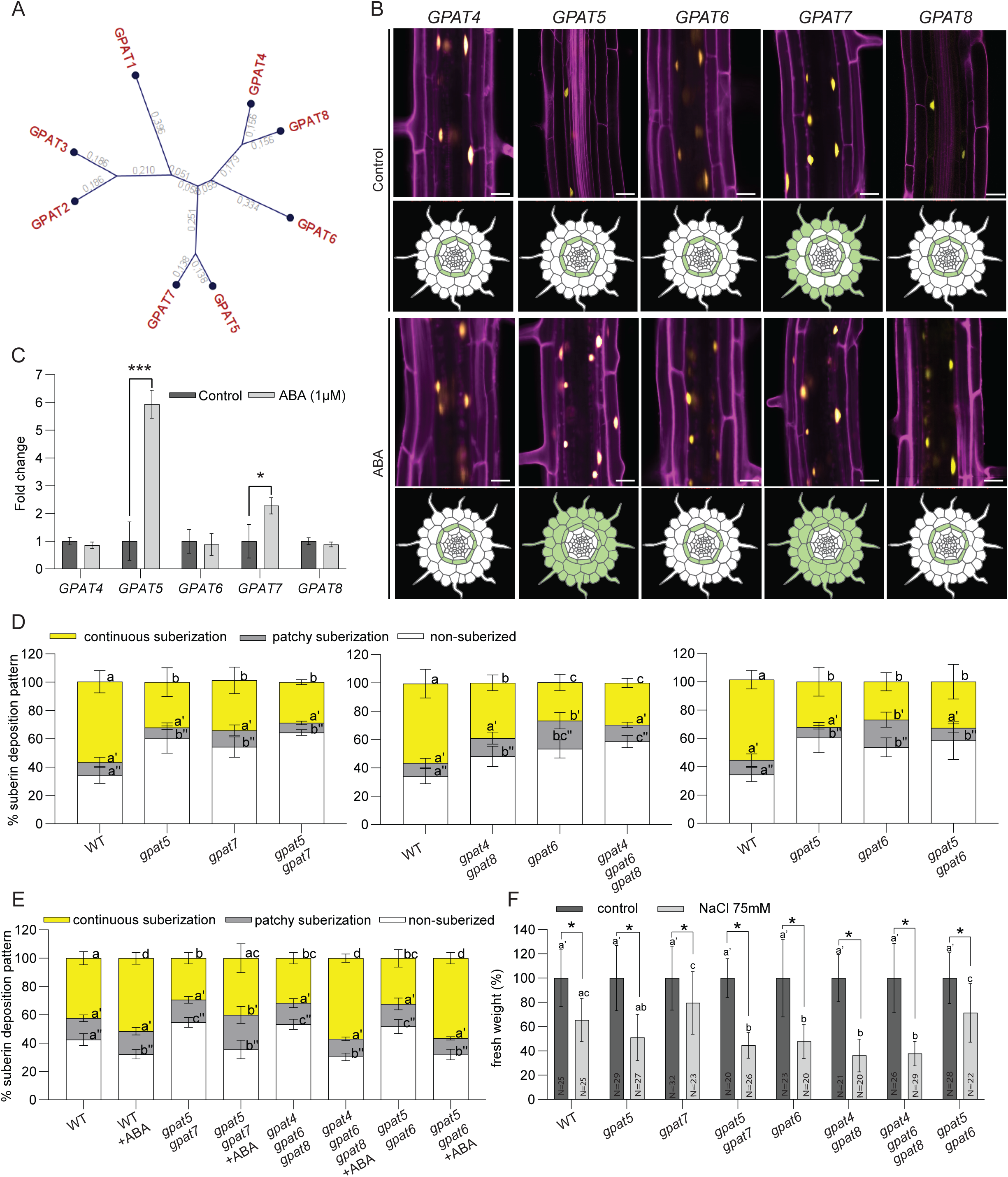
Relevance of the *GPAT4/6/8* and *GPAT5/7* clades for root suberization and salt stress resistance in seedlings. (A) Phylogenic relationship of land plant specific *GPATs* according to nucleotide sequence. Bootstrap analysis was based on 1,000 replicates (CLC workbench, QIAGEN). The number of base pair substitutions/sites is indicated at each branch. (B) *promoterGPATx:nls-GFP-GUS* reporter expression in transgenic Arabidopsis seedlings. GFP fluorescence is shown in yellow for each *GPAT* promoter. Propidium iodide (PI) fluorescence is presented in magenta as an indicator of root cell shape. A fully suberized root section closest to the hypocotyl was evaluated in 5-day-old roots that were grown for 4 days on half-strength MS medium and then transferred to half-strength MS medium supplemented with 1 µM abscisic acid (ABA) or methanol (control) for 20 h. 3D-Z projections of longitudinal views (upper panels) and schematic diagrams of cross-section views are presented (bottom panels). The cross-section views were adapted from an illustration of a root cross-section (https://doi.Org/10·6084/m9·figshare.c·3701038·v4)· See also Fig. S1. Scale bars, 25 µm. (C) Expression of *GPATs,* as evaluated by RT-qPCR in wild-type seedlings grown as described in B. Results are presented as the means of the fold-changes relative to the respective control + standard deviation (SD) of three biological replicates. Significant differences relative to the WT were determined by Student’s t-test. *, *p <* 0.05 and ***, *p <* 0.001. (D) Fluorol Yellow (FY) staining for suberin in the wild type (WT) and *gpat* mutants. Seedlings were grown for 5 days on half-strength MS medium under standard conditions. Quantification of the suberin pattern along the longitudinal axis of the root was performed. Data are means + SD, n ≥ 6. Different letters indicate significant differences, as determined by ANOVA with Tukey’s post-hoc test, *p <* 0.05. (E) Suberization of *GPA*T-clade mutants in response to abscisic acid (ABA), as evaluated by Fluorol Yellow (FY) staining. Entire roots of 5-day-old seedlings that were grown for 4 days on half-strength MS medium and then transferred to half strength MS supplemented with 1 µM ABA or methanol (control) for 20 h were evaluated. Different lowercase letters indicate significant differences determined by ANOVA with Tukey’s post hoc test, p-value < 0.05. (F) Fresh weight reduction in *gpat* mutants under salt stress. Plants were grown for 4 days on half-strength MS medium and then transferred to half-strength MS medium containing 75 mM NaCI for 9 days. Salt stress affected the growth of *gpat5 gpat7, gpat4 gpat8* and *gpat4 gpatβ gpat8* seedlings more than other genotypes. Mean of fresh weight + SD, n ≥ 20. Different lowercase letters indicate significant differences, as determined by ANOVA and Tukey’s post hoc test, *p-*value < 0.05. Significant differences between salt stress and control conditions are shown by an asterisk, as determined by Student’s t-test *: p < 0.05.

The five *GPAT*s expressed in the endodermis (*GPAT4–8*) were continuously transcribed along the roots of 5-day-old seedlings, when endodermal cells are fully differentiated. However, the initial position of *GPAT* expression along the root varied among these members. *GPAT5* expression began in the late differentiation zone, in agreement with the formation of suberin lamellae, as previously reported (30) (*SI Appendix*, Fig. S1*B*). Similarly, the expression of *GPAT6* and *GPAT7* began in the late differentiation zone, as evidenced by the block of propidium iodide diffusion into the endodermis, an indicator of a functional Casparian strip (*SI Appendix*, Fig. S1*B*) (32). By contrast, *GPAT4* and *GPAT8* were expressed in the elongation zone prior to Casparian strip formation (*SI Appendix*, Fig. S1*B*).

We also detected differences in the expression patterns of genes from both *GPAT* clades in response to exogenous ABA, which induces suberization, as previously reported (30). After ABA treatment, only *GPAT5* and *GPAT7* mRNA levels increased (Fig. 1*C*). In agreement with earlier reports (30), ABA treatment resulted in earlier *GPAT5* expression during root development in the endodermis and ectopic expression in cortical and epidermal cells (Fig. 1*B* and *SI Appendix*, Fig. S1*B*). *GPAT7* shared a similar expression pattern with *GPAT5*, whereas the expression patterns of *GPAT4*, *GPAT6*, and *GPAT8* did not change after ABA treatment (Fig. 1*B* and *SI Appendix*, Fig. S1*B*).

In summary, *GPAT*s of the *GPAT4*/*6*/*8* and *GPAT5*/*7* clades are expressed in the root endodermis following a specific developmental pattern. However, only the expression patterns of *GPAT5* and *GPAT7* changed in response to ABA treatment.

### *GPAT5*/*7* and *GPAT4*/*6*/*8* contribute to suberization and protect against mild salinity stress

To determine whether the GPATs of both clades act in suberin formation, we generated higher-order mutants by crossing the previously characterized single T-DNA insertion mutants *gpat5*, *gpat6*, and *gpat7* as well as the *gpat4 gpat8* double mutant (19, 25, 27, 29). We thus obtained mutants by knocking out an entire *GPAT* clade (the *gpat5 gpat7* double mutant and the *gpat4 gpat6 gpat8* triple mutant), together with the mixed-clade *gpat5 gpat6* double mutant. The *gpat5 gpat7* double mutant and *gpat4 gpat6 gpat8* triple mutant had a slightly stunted growth habit compared to the WT, while the *gpat5 gpat6* mutant grew as well as the WT (*SI Appendix*, Fig. S2*A* and *B*).

Since mutants with defects in suberization exhibit delayed suberin deposition (5, 30), we investigated the pattern of suberin deposition in the different *gpat* mutants by whole-mount staining of 5-day-old roots with fluorol yellow (FY), which stains lipid polyesters (33). In the WT, suberin deposition was patchy in younger parts of the roots and then became continuous, as observed earlier (30). All mutant genotypes showed a decreased extent of root suberization compared to the WT (Fig. 1*D*). In the *gpat5* and *gpat7* single mutants, endodermal suberization was strongly delayed, but the phenotype was not enhanced in the *gpat5 gpat7* double mutant (Fig. 1*D*). By contrast, *gpat4 gpat8* and *gpat6* had a less pronounced delay in suberization than the *gpat4 gpat6 gpat8* triple mutant (Fig. 1*D*). The *gpat5 gpat6* mutant did not have a stronger delay in suberization than either single mutant (Fig. 1*D*).

Notably, we observed the lower degree of suberization despite strong, potentially compensatory, upregulation of the remaining functional *GPAT* members in each mutant background (*SI Appendix*, Fig. S2*C–E*). In the *gpat5 gpat7* double mutant, *GPAT4, GPAT6*, and *GPAT8* expression increased 4-, 16-, and 2-fold, respectively, compared to the WT. In the *gpat4 gpat6 gpat8* triple mutant, *GPAT5* and *GPAT7* expression increased 15-and 8-fold, respectively. Exogenous ABA application further induced *GPAT5* and *GPAT7* expression in the mutants, suggesting that the upregulation of these genes in the mutants might occur independently of transcriptional regulation by ABA (*SI Appendix*, Fig. S2*C–E*).

We also investigated the suberization pattern along the root axis after ABA treatment. Suberization extended slightly toward younger root sections in the *gpat5 gpat7* mutant, with an expansion of the patchy zone, even when induction by ABA was only significant (and modest) for *GPAT8* transcript levels (Fig. 1*E* and *SI Appendix*, Fig. S2*C*). In *gpat4 gpat6 gpat8*, in which GPAT5 and GPAT7 are active and their encoding genes upregulated after ABA treatment, the suberin coverage was identical to the WT under the same conditions (Fig. 1*E* and *SI Appendix*, Fig. S2*D*). The *gpat5 gpat6* mutant also showed a WT-like suberization pattern after ABA treatment, likely due to the high and early induction of *GPAT7* (Fig. 1*E* and *SI Appendix*, Fig. S2*E*).

Suberin is well known for protecting plants against salinity stress via an ABA-dependent regulatory pathway (34, 35). Therefore, we examined the effect of mild salinity stress conditions, provided in the form of 75 mM NaCl treatment, on the growth of various *gpat* mutants after seedling growth had been established under normal conditions for 4 days. After 9 days of mild salinity stress, the *gpat5 gpat7* and *gpat4 gpat8* double mutants and the *gpat4 gpat6 gpat8* triple mutant exhibited diminished growth compared to WT seedlings, as determined by measuring their fresh weight (Fig. 1*F* and *SI Appendix*, Fig. S2*F*). Notably, the growth of the *gpat5 gpat6* mixed-clade mutant was similar to the WT under the same mild salinity-stress conditions.

In summary, members of both *GPAT* clades contribute to suberization and protection against mild salinity stress.

### *GPAT* clade mutants have characteristic changes in suberin amounts and composition

To investigate the roles of the different *GPAT* clade members in suberin formation, we analyzed the amounts and composition of ester-bound suberin components in 5-day-old seedlings, which have a suberized endodermis, and in 3-week-old plants, which have a suberized periderm in addition to a suberized endodermis (Fig. 2 and *SI Appendix*, Fig. S3) (36).

**Fig. 2.**
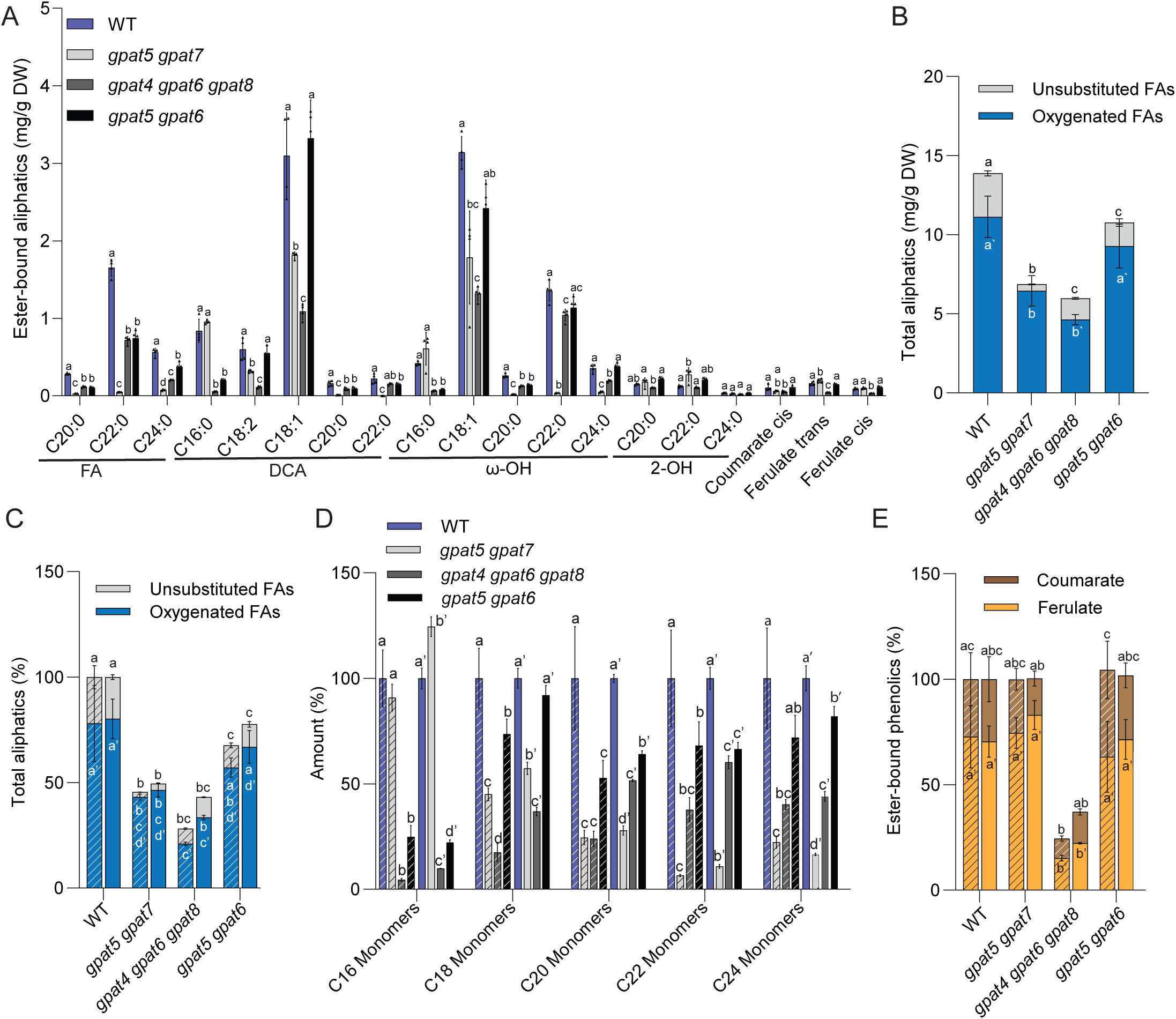
Characteristic alterations of suberin monomer composition in *GPAT* clade mutants. (A-B) Quantification of aliphatic and aromatic ester-bound suberin monomers isolated from roots of 3-week-old *GPAT* clade mutant plants in comparison to the wild type (WT). Suberin monomers (A) and the sum of the evaluated aliphatic compounds (total) grouped by substance classes (B) are presented. Plants were grown on half-strength MS medium. Values represent means ± SD, n = 4. Different lowercase letters indicate significant differences, as determined by ANOVA with Tukey’s post-hoc test, *p <* 0.05. Black triangles depict individual data points. FA, fatty acid; DCA; dicarboxylic acid; DW, dry weight. (C-E) Suberin monomers of roots of 5-days-old plants (stripped columns) and 3-weeks-old plants (uniform columns) of different genotypes: total aliphatic monomers groups by substance class (C), aliphatic suberin monomers grouped by chain-length (D), principal ester-bound hydroxycinnamic acids (E). Data are presented in % in comparison with the corresponding WT value. The values of 5-day-old WT plants represent a mean ± SD of all data presented in Fig. S3 (n=12). Values of 5-day-old mutant plants are based on data presented in Fig. S3 and of all 3-week-old plants in Fig. 2A (means ± SD, n = 4). Different lowercase letters indicate significant differences among genotypes (C, D) and among genotypes of the same developmental stage (D), as determined by ANOVA and Tukey’s post hoc test, p-value < 0.05. The complete data for 5-day-old plants are given in Fig. S3 and for 3-week-old plants in Fig. 2A.

At both developmental stages, *gpat5 gpat7* exhibited an approximately 50% drop in the amount of aliphatic suberin monomers (Fig. 2*A* and *C*; *SI Appendix*, Fig. S3*A*). The *gpat5* mutant showed decreased amounts of a wide range of suberin monomers, except for C16 monomers in 5-day-old seedlings. The amount and composition of very-long-chain monomers were less affected in the *gpat7* single mutant compared to the WT (*SI Appendix*, Fig. S3*A*). However, we observed reproducible, additive effects of *GPAT5* and *GPAT7* on C20 and C22 monomers (*SI Appendix*, Fig. S3*A*). The strong decrease in C22 monomers (80–90%) without any change in C16 monomers was characteristic of the suberin of the *gpat5 gpat7* double mutant at both developmental stages (Fig. 2*D* and *SI Appendix*, Fig. S3*A*).

The *gpat4 gpat6 gpat8* triple mutant showed a great reduction in the amounts of aliphatic suberin monomers of all chain lengths (Fig. 2*D* and *SI Appendix*, Fig. S3*B*), which resulted in a 75% and 55% drop in total aliphatic monomers in 5-day-old seedlings and 3-week-old plants, respectively (Fig. 2*B* and *D*). We observed an additive effect of *GPAT6* and *GPAT4/GPAT8* on most monomers, including C16 monomers, which reached only 5% of WT amounts in 5-day-old seedlings. This strong decrease in C16 monomer amounts was characteristic of the *gpat4 gpat6 gpat8* mutant at both developmental stages (Fig. 2*D* and *SI Appendix*, Fig. S3*B*).

We observed no additive effects in aliphatic suberin monomers in the *gpat5 gpat6* mixed-clade mutant, although 5-day-old seedlings of both single mutants showed characteristic reductions in the amounts of C16 and C22 monomers (*SI Appendix*, Fig. S3*C*). The overall suberin amount of *gpat5 gpat6* was 35% lower than that of the WT in 5-day-old seedlings and 22% lower in 3-week-old plants (Fig. 2*E* and *SI Appendix*, Fig. S3*C*). The abundance of numerous monomers decreased to a lesser extent in *gpat5 gpat6* than in the pure *GPAT* clade mutants at both stages of development (Fig. 2*A* and *SI Appendix*, Fig. S3*C*).

The amounts of ester-bound hydroxycinnamic acids also showed striking differences among *GPAT* clade mutants at both developmental stages (Fig. 2*E* and *SI Appendix*, Fig. S3*D– F*). The amounts of ferulate and coumarate were not significantly reduced in any *GPAT5*/*7* clade mutants nor in the *gpat5 gpat6* mixed-clade mutant in the roots of 5-day-old seedlings (Fig. 2*E*, *SI Appendix*, Fig. S3*D* and *F*). By contrast, all mutants of the *GPAT4*/*6*/*8* clade showed at least a 30% lower ferulate content than the WT, with the most drastic drop (70–80%) seen in the *gpat4 gpat6 gpat8* triple mutant (Fig. 2*E* and *SI Appendix*, Fig. S3*E*).

In summary, knocking out the genes from each *GPAT* clade led to characteristic changes in suberin amount and composition, indicating that these genes play redundant roles within each clade and non-redundant roles across the different clades during suberin formation in the endodermis and periderm.

### Knockout of the *GPAT5*/*GPAT7* clade leads to the deposition of globular amorphous suberin

We examined the ultrastructure of the suberized cell wall in the different *GPAT* clade mutants by transmission electron microscopy (TEM) in the endodermis of 5-day-old seedlings (Fig. 3*A* and *SI Appendix*, Fig. S4) and phellem cells of 5-week-old plants, comprising some of the mature periderm (Fig. 3*B* and *SI Appendix*, Fig. S4). In the WT, we observed a layer of suberin with a typically lamellated substructure at both developmental stages. However, all mutants showed characteristic ultrastructural differences (Fig. 3 and *SI Appendix*, Fig. S4). We detected a very thin but continuous layer of suberin in the *gpat4 gpat6 gpat8* mutant in both endodermal and phellem cells with a lamellated substructure (Fig. 3 and *SI Appendix*, Fig. S4).

**Fig. 3.**
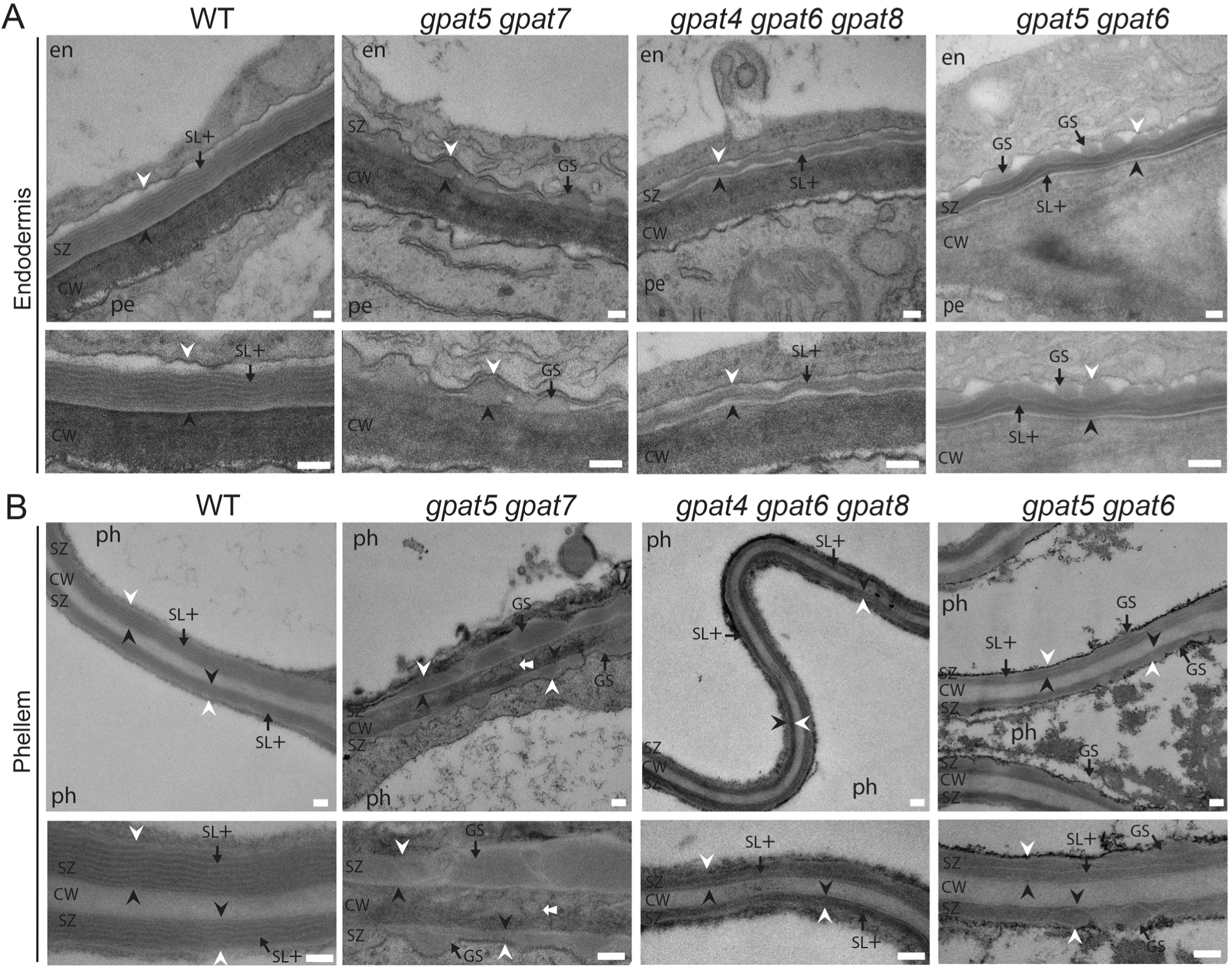
Characteristic alterations of the suberin ultrastructure in *GPAT* clade mutants. (A-B) Transmission electron micrographs (TEM) of the suberized cell wall in endodermal (A) and phellem (B) cells in *GPAT* clade mutants compared to the wild type (WT). The suberized endodermis was evaluated on the side neighboring the pericycle in 5-days-old seedlings grown on half-strength MS medium (A). Phellem cells of 5-week-old soil-grown plants having a mature periderm were analyzed (B). Higher magnifications are shown in the lower panels. See also Fig. S4 for additional pictures with different magnifications. Scale bars, 100 nm. CW, cell wall; SZ, suberin zone. White arrowhead, plasma membrane; black arrowhead, junction between suberin zone (SZ) and polysaccharide cell wall (CW); Black arrows points at polyester deposits of different structures: GS, globular suberin; SL+, suberin layer with a lamellated substructure; en, endodermal cell; ep, epidermal cell; ph, phellem cell, pe, pericycle cell. White arrow points at globular suberin embedded in the polysaccharide cell wall.

Strikingly, we did not observe a continuous lamellated suberin layer in the *gpat5 gpat7* mutant. In its place, we noticed globular suberin deposits that were irregular in size and amorphous in structure, leaving gaps in the suberin coverage of the cells (Fig. 3 and *SI Appendix*, Fig. S4). In phellem cells, amorphous globular structures were also visible within the polysaccharide cell wall, indicating an overall highly modified structure of the cell wall continuum.

The polyester deposits in the *GPAT4*/*6*/*8* and *GPAT5*/*7* clade mutants displayed distinct features. In agreement with this finding, we observed both types of polymer deposits in the endodermal and phellem cells of the *gpat5 gpat6* mutant. Indeed, we detected a continuous layer of suberin possessing an irregular lamellae substructure and globular structures on the side of the suberin layer, oriented toward the plasma membrane (Fig. 3 and *SI Appendix*, Fig. S4).

We assessed the degree of suberization in *GPAT* clade mutants in 5-day-old seedlings after ABA treatment (30). All genotypes except the *gpat5 gpat7* double mutant showed a strong increase in suberization after ABA treatment, as evaluated by FY staining (Fig. 1*E* and *SI Appendix*, Fig. S5*A*). In addition to increased suberization in the endodermis after ABA treatment, a newly formed suberizing zone with little substructure was present in the cortical cells of WT seedlings (*SI Appendix*, Fig. S5*C–E*), as previously reported (30). The suberizing zone in the endodermis increased in thickness after ABA treatment in the *gpat4 gpat6 gpat8* triple mutant, even when the number of suberin lamellae did not increase. In addition, a newly formed suberizing zone was observed in the cortical cells of *gpat4 gpat6 gpat8*, like in the WT (*SI Appendix*, Fig. S5*C–E*).

In *gpat5 gpat7*, endodermal suberin also showed characteristic globular structures after ABA treatment. Furthermore, we observed only limited deposition of globular suberin in the cortex, which did not form a continuous layer (*SI Appendix*, Fig. S5*C* and *E*). This result is in agreement with the finding that only a small amount of GPAT activity was derived from ABA-inducible *GPAT8* in the *gpat5 gpat7* double mutant (*SI Appendix*, Fig. S2*C*).

In the *gpat5 gpat6* mixed-clade mutant, the suberized zone of the endodermis did not expand significantly with ABA treatment (*SI Appendix*, Fig. S5*B* and *D*), but we observed no globular suberin structures, in contrast to the control condition in the absence of ABA treatment, indicating that strong *GPAT7* induction by ABA may lead to fewer globular deposits (*SI Appendix*, Figs. S2*E* and S5*B*). In the cortex, an amorphous suberizing zone with little substructure was present, similar to that observed in the WT (*SI Appendix*, Fig. S5*C* and *E*).

In summary, we observed characteristic changes in suberin ultrastructure in the different *GPAT* clade mutants, pointing to distinct contributions of each *GPAT* clade to suberin formation. In particular, we observed a striking suberin ultrastructure in the *gpat5 gpat7* mutant, in which the continuous suberin layer with its characteristic lamellae was lost and replaced by distinct globular, amorphous suberin deposits.

### An active phosphatase domain of GPAT4, GPAT6, and GPAT8 is required for suberin and cutin deposition

We assessed whether the presence of an active phosphatase domain plays a significant, positive role in suberin formation. To this end, we targeted crucial amino acids in the phosphatase-active sites of GPAT4, GPAT6, and GPAT8 by site-directed mutagenesis (22). In each GPAT, we replaced the two conserved aspartic acid (D) residues in motif III of the phosphatase domain with lysine (K) residues, as previously described (*SI Appendix*, Fig. S6*A*) (23, 37). We then introduced a construct expressing the unmutated or mutated version of *GPAT4* (*GPAT4*\*) driven by its native promoter (*pGPAT4:GPAT4* or *pGPAT4:GPAT4\**) as well as the intact or mutated version of *GPAT8* (*GPAT8*\*) driven by its native promoter (*pGPAT8:GPAT8* or *pGPAT8:GPAT8*\*) into the *gpat4 gpat8* mutant. We generated comparable constructs for *GPAT6* (unmutated *GPAT6* and mutated *GPAT6* [*GPAT6\**]) and introduced them into the *gpat6* mutant. We selected transgenic lines in which the transgenes were expressed at levels similar to or even higher than the native genes in the WT (*SI Appendix*, Fig. S6*B*).

The suberin deposition pattern of the root, as evaluated by FY staining, was not rescued by introducing any *GPAT*\* construct encoding a GPAT with a mutated phosphatase domain, in contrast to the full complementation observed with the unmutated *GPAT* constructs (*SI Appendix*, Fig. S6*C*). However, chemical analysis of suberin amount and composition revealed that the *pGPAT4:GPAT4*\* transgene partially rescued the overall amount of suberin monomers in the *gpat4 gpat8* double mutant (Fig. 4*A* and *SI Appendix*, Fig. S7*A* and *C*). In agreement with this finding, the suberin layer (as observed by TEM) was thicker in *gpat4 gpat8* harboring *pGPAT4:GPAT4\** than in *gpat4 gpat8*, even though it did not reach the thickness seen with the *pGPAT4:GPAT4* transgene or in the WT (*SI Appendix*, Fig. S8*D* and *E*). The introduction of *pGPAT8:GPAT8\** in the *gpat4 gpat8* double mutant and *pGPAT6:GPAT6\** in the *gpat6* mutant also partially rescued the defect in suberization, with some variability among experimental replicates (Fig. 4*B* and *C*, *SI Appendix*, Figs. S7*B* and *C*, S8*B*, *C*, *E*, *F*).

**Fig. 4.**
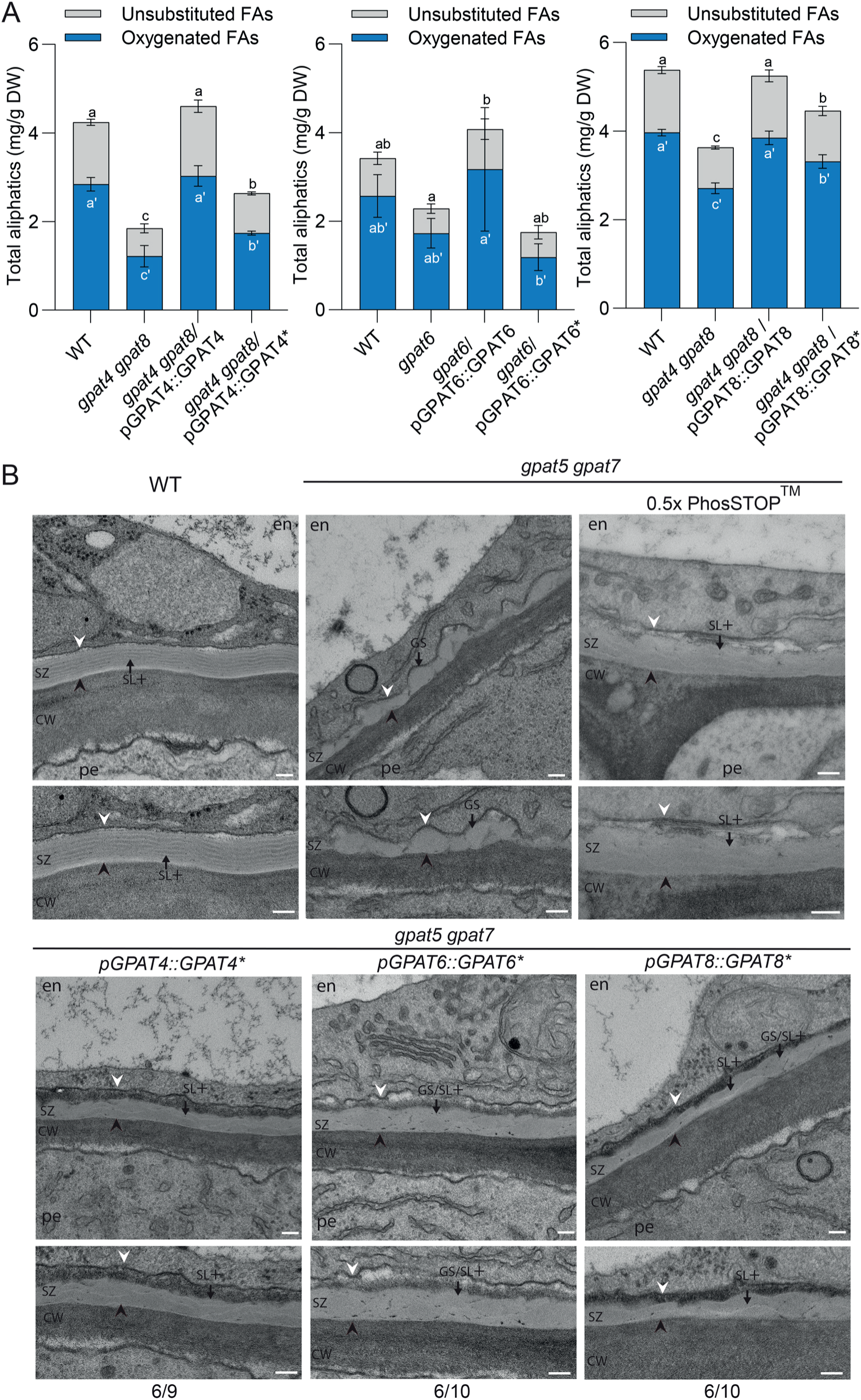
GPATs with and without phosphatase activity play complementary roles in suberin formation. (A-B) Modifications of phosphatase activities in *GPAT* clade mutants. (A) Effect of GPAT phosphatase active site mutations on suberin formation (GPAT4*, left; GPAT6*, middle; GPAT8*; right). The amount of aliphatic suberin components were evaluated in *GPAT* clade mutants transformed with the respective WT gene (GPAT4, GPAT6 or GPAT8) as well as the mutated gene, encoding for GPATs, in which the two conserved aspartic acid residues of the phosphatase domain were mutated to lysine residues (GPAT4*, GPAT6* or GPAT8*). Only transgenic lines with an expression level similar or higher than that of the WT were analyzed (Fig. S6B). Plants were grown on half-strength MS medium for 5 days. Values represent the means +/-SD, n = 4. Different lowercase letters indicate significant differences among the genotypes, as determined by ANOVA with Tukey’s post hoc test, p-value < 0.05; FA, fatty acid; DW, dry weight. See also Fig. S7 for complete datasets on suberin formation and Fig. S9 and S10 on cutin formation. (B) Effect of different strategies interfering with GPAT phosphatase activities on the suberin ultrastructure in the *GPAT5/7* clade mutant. Transmission electron micrographs (TEM) of the suberized endodermis in *gpatδ gpat7* that were either treated with a phosphatase inhibitor PhosSTOP™ (0.5x) or expressed a phosphatase-dead GPAT compared to the wild type (WT) and the gpatδ gpatľ double mutant. The suberized endodermis was evaluated on the side neighboring the pericycle. Plants were grown for 5 days on half-strength MS medium, except these treated with phosphatase inhibitor that were grown for 3 days on half-strength MS medium and for 2 days on half-strength MS supplemented with the phosphatase inhibitor PhosSTOP™ (0.5x) (See also Fig. S12 and S13 for controls and additional pictures). Transgenic *gpatδ gpat7* plants of the T1 generation expressing a mutated *GPAT* gene encoding for a member of the *GPAT4/6/8* clade, in which the two conserved aspartic acid residues of the phosphatase domain were mutated to lysine residues and expressed under their respective native promoter *(proGPAT4:GPAT4*, proGPAT6:GPAT6*, proGPAT8:GPAT8*)* were evaluated. Higher magnifications are shown in the lower panels (see also Fig. S14 for additional examples). The numbers underneath the pictures indicate how many roots out of all analyzed T1 plants showed an improved suberin organization, i.e. a continuous suberin layer, in the majority of endodermal cells. CW, cell wall; SZ, suberin zone. White arrowhead, plasma membrane; black arrowhead, junction between suberin zone (SZ) and polysaccharide cell wall (CW). Arrows point to polyester deposits of different structures; GS: globular suberin; SL+: suberin layer with a strongly lamellated substructure; GS/SL+: intermediate structure with globular suberin as well as lamellated substructure. Scale bar, 100 µm.

We also investigated the effects of mutating the phosphatase domains of GPAT4, GPAT6, and GPAT8 on cutin formation. Transgenic expression of *GPAT4* or *GPAT8* variants carrying a mutated version of the phosphatase domain driven by the native promoter led to a partial rescue of the strong reduction in cutin amounts observed in *gpat4 gpat8* rosette leaves (*SI Appendix*, Fig. S9*A* and *B*). Notably, the barrier properties of the leaf cuticle in the *gpat4 gpat8* double mutant were fully restored to WT levels by the expression of *GPAT4*\* or *GPAT8*\* (*SI Appendix*, Fig. S9*C*). The markedly lower amount of floral cutin in the g*pat6* mutant as well as the barrier properties of the petal cuticle were only partially rescued by *GPAT6*\* but fully rescued by *GPAT6* (*SI Appendix*, Fig. S10*A* and *B*). In summary, GPATs containing an active site mutation in the phosphatase domain failed to fully complement the defects in suberin or cutin deposition and associated phenotypes in the mutants. The importance of the active phosphatase domain in GPAT4, GPAT6, and GPAT8 can vary among organs and the type of polyester.

Due to their active phosphatase domains, GPAT4, GPAT6, and GPAT8 form *sn2*-MAGs, in contrast to GPAT5 and GPAT7, which form LPAs (19, 23). We investigated root waxes extracted by washing roots from 3-week-old WT and mutant plants with chloroform to gain insights into the amounts and composition of suberin precursors, with a focus on MAGs and LPAs (*SI Appendix*, Fig. S10*A*). Root waxes contained C24:0-*sn2*-MAG as their principal component (*SI Appendix*, Fig. S11*A*) and MAGs with other chain lengths of the acyl moiety, but no LPAs (*SI Appendix*, Fig. S11*B*). We only detected MAGs with unsubstituted acyl chains, as reported earlier (38, 39). Neither MAGs nor LPAs with a hydroxylated acyl moiety were detected, suggesting that suberin precursors were below the detection limit in root waxes (*SI Appendix*, Fig. S11).

### GPATs lacking phosphatase activity are crucial for the formation of suberin lamellae

GPAT5/7 are characterized by the presence of an inactive phosphatase domain. We tested whether the absence of phosphatase activity is required to generate suberin lamellae by treating WT and *gpat5 gpat7* seedlings with the phosphatase inhibitor PhosSTOP^TM^, with *gpat5 gpat7* retaining only GPATs with an active phosphatase domain. Since PhosSTOP^TM^ decreases but does not eliminate the activities of different classes of phosphatases in plants (Roche), PhosSTOP^TM^ treatment was not detrimental to the seedlings, although the growth of both WT and the *gpat5 gpat7* seedlings was inhibited (*SI Appendix*, Fig. S12*A* and *B*).

The suberization pattern during root development, the thickness of the suberin layer, and the ultrastructure of the lamellated suberin were unaffected by PhosSTOP^TM^ treatment in WT seedlings (*SI Appendix*, Fig. S12*C*). By contrast, treating *gpat5 gpat7* with the phosphatase inhibitor led to a strong reduction in globular suberin deposits with an amorphous appearance; instead, a continuous suberin layer with an irregularly lamellated substructure formed (Fig. 4*B*, *SI Appendix*, Fig. S12*C*, Fig. S13). Examination of the cell wall by electron tomography revealed that these irregular lamellae, which could often only be visualized over short distances were indeed continuous lamellae (*SI Appendix*, Fig. S14 and Supplementary Movies S1–S8).

To investigate whether an inactive phosphatase domain in GPATs is required for suberin lamellae formation, we introduced the transgenes *pGPAT4:GPAT4*, pGPAT6:GPAT6*\*, or *pGPAT8:GPAT8*\* encoding phosphatase-dead versions of GPAT4 (GPAT4*), GPAT6 (GPAT6*), or GPAT8 (GPAT8*) driven by their respective native promoters into the *gpat5 gpat7* double mutant. We investigated suberin ultrastructure in the endodermis of 5-day-old seedlings of the first generation after transformation (T1). In contrast to the distinct globular suberin deposits of variable size seen in *gpat5 gpat7*, we observed a continuous layer of suberin with an irregular lamellated structure in several independent transgenic *gpat5 gpat7* seedlings producing phosphatase-dead versions of GPATs (Fig. 4*B*, *SI Appendix*, Fig. S15). In particular, in transgenic *gpat5 gpat7* seedlings expressing *GPAT4*\* and occasionally in those expressing *GPAT8*\*, the suberin layers had a lamellated substructure, in which the dark suberin lamellae of irregular dimensions alternated with thin electron-lucent lamellae with an appearance similar to that in the WT (Fig. 4*B*, *SI Appendix*, Fig. S15). Expression of all *GPAT*\* variants also led to less contrasted suberin, with partially globular deposits within the lamellae of various dimensions, as well as electron-dense inclusions, as shown for *GPAT6* in Fig. 4*B* and for other genotypes in *SI Appendix*, Fig. S15.

In summary, pharmacological inhibition of endogenous phosphatases and expression of various phosphatase-dead variants of GPATs both led to strikingly similar suberin structures, characterized by a continuous suberin layer with lamellae, even when irregular, indicating that the loss of phosphatase activity in GPATs plays a crucial role in suberin lamellae formation.

## Discussion

### *GPAT4/6/8* and *GPAT5/7* play non-redundant roles in suberin formation

We demonstrated that the *GPAT4*/*6*/*8* and *GPAT5/7* clades contribute to suberin formation by characterizing suberin deposition in the roots of several *GPAT* clade mutants. Both mutants with knock-out of an entire *GPAT* clade (*gpat5 gpat7* and *gpat4 gpat6 gpat8*) showed a strong decrease in suberin content and thickness during endodermal suberization and periderm formation, indicating that these clades play non-redundant roles throughout root development (Fig. 2*B*, *C* and *E*). Growth defects in these pure *GPAT* clade mutants were associated with reduced suberin amounts, in agreement with the important functions of suberin in water relations and stress tolerance (40). The finding that GPATs from different clades function independently in suberin formation is supported by the more limited drop in suberin amount and the normal growth seen in the mixed-clade *gpat5 gpat6* double mutant (Fig. 1*F*, Fig. 2, *SI Appendix*, Fig. S2*A* and *B*).

### GPAT4, GPAT6, and GPAT8 contribute broadly to polyester formation

Suberin monomers of all chain lengths accumulated to lower amounts in the roots of 5-day-old *gpat4 gpat8* double mutant seedlings than the WT, indicating no chain length preference for GPAT4 or GPAT8 (*SI Appendix*, Fig. S3*B*). By contrast, cutin in the leaf, stem, and root cap of the *gpat4 gpat8* mutant was predominantly deficient in C16 and C18 dicarboxylic acids (*SI Appendix*, Fig. S8*A* and *B*) (25, 33). Similarly, the floral cutin of *gpat6* was predominantly deficient in C16 monomers (*SI Appendix*, Fig. S9*A*) (27, 41), while *gpat6* was affected in all suberin monomers (*SI Appendix*, Fig. S3*B*). Nevertheless, the amounts of C16 suberin monomers were markedly lower in *gpat6* (70–75% reduction) compared to monomers of other chain lengths (40–45% reduction) (*SI Appendix*, Fig. S3*B*), as observed recently (28). These findings suggest a possible chain length preference for C16 monomers by GPAT6. The stronger effect seen in the amounts of all suberin monomers in *gpat4 gpat6 gpat8* demonstrates the concerted action of GPAT4/6/8 clade members in suberin formation.

### An active phosphatase domain in GPAT4/6/8 clade members plays a role in suberization

GPAT4/6/8 clade members carry an active phosphatase domain and therefore synthesize monoacylglycerols (MAGs), which distinguishes them from GPAT5/7 clade members (19, 23). Complementation of the respective mutants with *GPAT4*\*, *GPAT6*\*, and *GPAT8*\*, producing protein variants with a mutated phosphatase domain active site, did not fully rescue the amount of polyester for cutin or suberin biosynthesis in the mutant (Fig. 4*A*, *SI Appendix*, Figs. S6–S10). These results highlight the importance of an active phosphatase domain in enzymes encoded by the *GPAT4/6/8* clade. The reason for certain differences in the requirements of the active phosphatase domain for the biosynthesis of suberin and cutin in different organs of Arabidopsis still needs to be investigated.

### GPAT5 and GPAT7 are adapted for suberin formation

*GPAT5* expression is tightly linked to suberin biosynthesis (29, 30). As previously observed, the *gpat5* mutant displayed a strong decrease in the amounts of C22 to C26 very long acyl chains (*SI Appendix*, Fig. S3*A*) (29). In addition, we revealed that *GPAT7*, encoding a protein with 88% amino acid identity to GPAT5, also contributes to root suberin formation. Neither *GPAT5/7* clade members affect the incorporation of C16 monomers into suberin. Instead, they affect the incorporation of C18 and longer monomers (Fig. 2*A* and *SI Appendix*, Fig. S3*A*), which are typical suberin components (15).

The expression of *GPAT5/7* clade members, but not the *GPAT4/6/8* clade, was strongly induced by ABA treatment under our growth conditions (Fig. 1*C* and *SI Appendix*, Fig. S1). Considering the high expression level of *GPAT7* in the *gpat5 gpat6* double mutant indicating compensation reactions (*SI Appendix*, Fig. S2*D* and *E*), *GPAT7* might also have partially compensated for the loss of GPAT5 in previous experiments (29). Both *GPAT5*/*7* clade members showed a dominant role in ABA-dependent suberin formation in the cortex (*SI Appendix*, Fig. S5*C*). In contrast to our observations, Shukla et al. did not observe differences in transcript levels between the *GPAT5/7* and *GPAT4/6/8* clade members after the external application of ABA pointing to differences in the regulatory circuits under different environmental conditions (42).

### GPATs without phosphatase activity are required for suberin lamellation

In *gpat5 gpat7*, only distinct amorphous globular structures were deposited, indicating that an important element was missing that was needed to structure suberin into lamellae (Fig. 3, *SI Appendix*, Fig. S4). GPAT5 and GPAT7 have an inactive phosphatase domain due to the presence of mutations in their active sites, suggesting that the absence of phosphatase activity might be important for the formation of suberin lamellae. Supporting this idea, general inhibition of phosphatases by pharmacological treatment led to the formation of suberin with a lamella substructure (Fig. 4*B*, *SI Appendix*, Figs. S12, S13; Supplementary Movie S8). The requirement of an inactive phosphatase domain in GPATs was further supported by introducing a transgene producing phosphatase-dead variants of GPAT4, GPAT6, or GPAT8 into the *gpat5 gpat7* double mutant, leading to the formation of a continuous suberin layer, including lamellae, although irregular in shape (Fig. 4B, *SI Appendix*, Fig. S15). These suberin lamellae formed in the absence of GPAT5/7 acyltransferase activity, highlighting the importance of the inactive phosphatase domain in GPATs. These observations suggest the importance of LPAs in suberin formation. Clearly, both GPAT clades are required for forming stacks of highly organized suberin lamellae that are typical for seed plants (Fig. 5) (43). Additional and yet unknown requirements for the formation of suberin lamellae must exist, since despite the presence of GPAT5/7 in the cortex, the suberin that formed was present in a continuous layer without lamellated structures (*SI Appendix*, Fig. S5*C*).

**Fig. 5.**
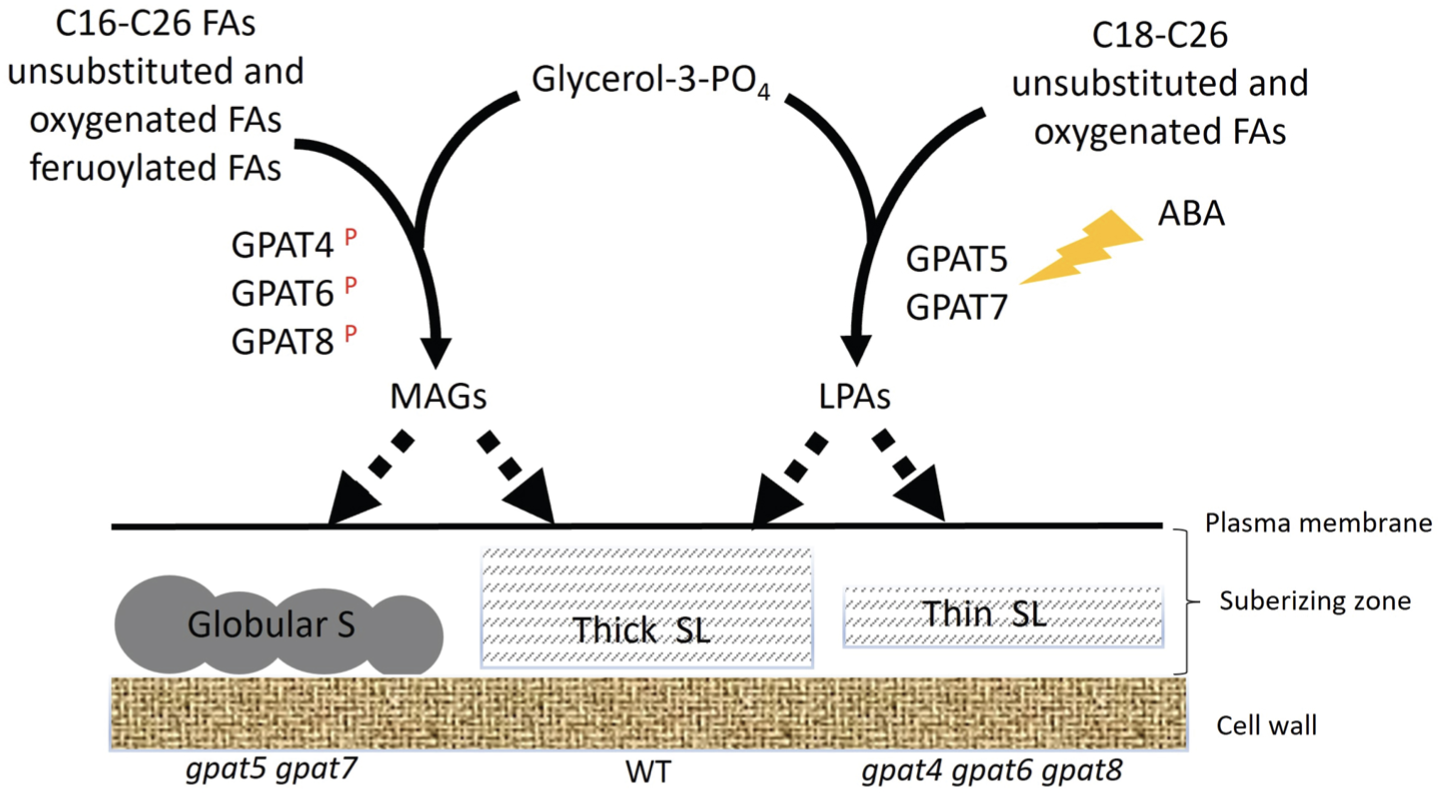
Schematic diagram summarizing the role of GPATs in suberin formation. GPATs with an active phosphatase domain (P) synthesize monoacylglycerols (MAGs) from a wide range of substrates lead to globular suberin formation when active alone. ABA-inducible GPATs lacking an active phosphatase domain synthesize lysophosphatidic acids (LPAs) and form a thin lamellated suberin layer when active alone. Both *GPAT* branches contribute to optimal suberin lamellae formation.

MAGs and LPAs have different physical properties, potentially requiring different export mechanisms towards the apoplast. During cutin formation, MAGs are exported by ATP-binding cassette G (ABCG) transporters (44). Similarly, suberin formation has been associated with ABCG transporter functions as well as extracellular vesicular bodies (EVBs), which are particularly abundant in suberizing cortical cells (45–48). Since cortex suberization highly depends on GPAT5/7 (*SI Appendix*, Fig. S5*C*), it is tempting to speculate that in roots, suberin precursors generated by GPAT5/GPAT7 are transported via EVBs (49).

Whether LPAs are required for the export of suberin precursors essential for the lamellated structures of suberin or for their assembly in the apoplast remain challenging questions for future research.

### Evolution of plant polyesters with specialized functions

GPATs of the GPAT4/6/8 clade are required for the formation of cutin and sporopollenin, which are thought to be the most ancient plant polyesters (50). *GPAT6* homologs are present in early plant lineages, which is consistent with the finding that GPATs carrying an active phosphatase domain are ancestral (19, 24).

Recent studies have shown that suberin lamellae have evolved in seed plants, contributing to higher drought tolerance in seed plants by conferring improved water transport properties (43). This evolution was accompanied by a burst of gene duplications and subsequent functional specialization of suberin biosynthesis genes (43). Our findings provide detailed insights into the roles of GPATs in suberization and reveal a specific role for the GPAT5/7 clade in the formation of suberin lamellae. The diversification of the GPAT family during evolution likely played an essential role in plant adaptation to different environmental conditions.

## Materials and Methods

### Plant materials

For all experiments, *Arabidopsis thaliana* (accession Columbia-0 [Col-0]) was used. Mutants were already described: *gpat4*, *gpat8*, *gpat4 gpat8* (25), *gpat5-1* (29), *gpat6-2* (27) and *gpat7-3* (19). Double and triple mutants were obtained by crossing the respective single mutants. *proGPAT5:nls-GFP-GUS* transcriptional reporter line was previously described (31).

### Growth conditions

For all experiments, plants were germinated on half-strength Murashige and Skoog (MS) medium with 0.7% (w/v) agar (Duchefa) (solid half-strength MS medium). Sucrose (0.3%, w/v) was regularly added for growth of *gpat4 gpat8* as well as *gpat4 gpat6 gpat8* seedlings, as well as respective controls. While the *gpat5 gpat6* and *gpat5 gpat7* double mutants germinated and grew as well as the WT on half-strength MS plates without sucrose, the *gpat4 gpat8* and *gpat4 gpat6 gpat8* mutants required sucrose for germination and growth on half-strength MS plates. After verifying that a small amount of sucrose (0.3%) did not affect suberin deposition (*SI Appendix*, Fig. S16*A* and *B*), we performed subsequent experiments growing these genotypes in the presence of 0.3% (w/v) sucrose. Seeds were surface sterilized, sown on plates, incubated for 2 days at 4 °C for stratification, and grown vertically in growth chambers at 22 °C under continuous light (100 μE). For transformation, seed amplification as well as for experiments on flowers, plants were grown on soil under long day light conditions (16-h light/8-h dark). Analysis of the leaf cuticle was performed with plants grown on soil under short day conditions (8-h light/16-h dark) for five weeks.

### Phytohormonal and phosphatase inhibition treatments

ABA was stored as a 50 mM stock solution in methanol. Seedlings were transferred to solid half-strength MS medium supplemented with 1 μM ABA or methanol (control), the transfer was done when the seedlings were 4 days old and were harvested after 20 h. For visualization of suberin lamellation in the cortex, seedlings were transferred after 4 days and harvested after 48 h. For salt experiments, the seedlings were grown on solid half-strength MS medium for 4 days and transferred to medium supplemented with 75 mM NaCl for 9 days before harvesting. For phosphatase inhibitor experiments, one tablet of the phosphatase inhibitor PhosSTOP^TM^, Roche Switzerland) was solved in 1 ml water to obtain the 10X solution and diluted to the indicated concentration in solid half-strength MS medium. Seedlings were transferred from half-strength MS medium to phosphatase inhibitor medium after 3 days and harvested after additional 2 days.

### Generation of transgenic lines

To generate pENTRY L4-pGPAT-R1, between 1.7-kb and 2.2-kb fragments upstream of each *GPAT* were amplified and cloned into pDONR P4-P1 using KpnI and XbaI restriction site. All transcriptional reporter lines were generated by recombining the corresponding pENTRY L4-pGPAT-R1 entry clone with pENTRY L1-nlsGFP-GUS-L2 (33) into the pFR7m24GW including a FastRed cassettes for transgenic seed selection (51) using the Gateway Technology (Lifesciences). Site-directed mutagenesis transgenic plants were generated by using the Q5 ^®^ Site-Directed Mutagenesis Kit (New England Biolabs) in the pDONR221 vector and recombined with the above-mentioned promotor constructs into the pFR7m24GW vector using the Gateway Technology. All constructs were transformed by heat shock into *Agrobacterium tumefaciens* GV3101 strain and then transformed into plants by floral dipping (52).

### Histological staining

For quantification of suberin deposition, five-day-old roots were stained using Fluorol Yellow (FY 088, Santa Cruz Biotechnology). Seedlings were incubated in methanol at room temperature for at least 3 days, stained with FY 088 (0.01% [w/v], in methanol) for 1 h at room temperature, rinsed in methanol and counterstained with aniline blue (0.5% [w/v] in methanol) at room temperature for 1 h in darkness and washed. For suberin staining, a stock solution of 1% (w/v) FY in DMSO was prepared as described previously (53). A working staining solution of 0.01% (w/v) FY was prepared by diluting the DMSO-based FY stock directly in Clearsee. Roots were stained for 2 h, washed in Clearsee solution for 10 min and visualized using 1-well chambered cover glass (ThermoFisher Scientific, catalog no. 155361).

### Permeability assay

To assess cuticle permeability 10–15 fully expanded flowers as were submerged in 500 mg l^-1^ Toluidine Blue (TB) 0.001% (v/v) Tween 20 solution under gentle agitation for 1.5 h and then rinsed thoroughly with water. Leaf 5-6 and leaf 8-9 were stained with a single drop (10 µl) of the above-described toluidine solution for 1 h and rinsed with water (54).

### Chemical analysis of polyesters

Chemical analysis of suberin was performed on roots of 5-day-old seedlings (30, 42). 200 mg of seeds were grown on nylon mesh (200 mm pore size). Chemical analysis of suberin of 3-week-old roots was performed on roots of 6 plants per replicate. Analysis of leaf cutin was performed on one 5-weeks-old entire rosette per replicate; flower cutin on 20 fully open flowers per replicate (15). The amounts of unsubstituted C16 and C18 fatty acids were not evaluated because of their omnipresence in the plant and in the environment.

For the analysis of root waxes, roots from 3-week-old plants were harvested and immediately immersed in 7 ml hot chloroform and shacked for 1 min. Monoheptadecanoin (17:0 monoacylglycerol) (Sigma) and Tetracosane (C24 alkane) (Sigma) were added as internal standards. The samples were split, and half of the root wax extraction was silylated with 50 µl, each, of pyridine and N,O-bis(trimethylsilyl)trifluoroacetamide at 70°C for 1 h. Derivatized samples were evaporated to dryness (without heating) and dissolved in hexane. Derivatized samples were separated on an Agilent GC-MS/FID system equipped with two Agilent 5MS capillary columns using the following temperature profile: starting at 90°C. heating with 40°/min to 220°C, with 2°C/min to 320°C, holding for 10 min. The remaining root wax extracts were analyzed by targeted LC-MS/MS analysis as described before (55). A brief summary of the procedure including modifications: the wax extracts were dried under streaming nitrogen and then extracted with propan-2-ol:hexane:water 60:26:14 (v/v/v). After drying under streaming nitrogen, the samples were dissolved in 200 µl of tetrahydrofuran:methanol:water 4:4:1 (v/v/v) (TMW). The samples were divided, and half of the samples were methylated with trimethylsilyldiazomethane for LPA analysis. The methylated and the non-methylated proportion of the samples were dried under streaming nitrogen and each dissolved in 45 µl TMW. For chromatographic separation, the flow rate was 0.13 min/min, starting with 40% of solvent B. For tandem mass spectrometric detection in multiple reaction monitoring mode, the target precursor ions were [M+NH_4_]+ for MAGs and [M+Me-H]^-^for LPA analysis. The target mass transitions were used to diagnose the acyl chain composition of the molecular species by using the different productions of either [FA-H_2_O+H]^+^ in positive ion mode (MAG) or [FA-H]^-^in negative ion mode (LPA).

### Microscopy techniques

Confocal laser-scanning microscopy images were obtained using either a Zeiss LSM 880 (with Zen 2.1 SP3 Black edition), and Leica LSM700. For fluorophores, the following excitation and detection windows were used. GFP, fluorol yellow: 488 nm, 500–530 nm, propidium iodide: 520 nm, 590–650 nm.

For transmission electron microscopy samples were preserved either by chemical fixation or high-pressure freezing, as described in detail in the Supplemental methods. Micrographs and panoramic were taken with a transmission electron microscope FEI CM100 (FEI) at an acceleration voltage of 80 kV with a TVIPS TemCamF416 digital camera (TVIPS) using the software EM-MENU 4.0 (TVIPS). Panoramas were aligned with the software IMOD (56). For electron tomography, the area of interest was taken with a transmission electron microscope JEOL JEM-2100Plus (JEOL Ltd., Akishima, Tokyo, Japan) at an acceleration voltage of 200 kV with a TVIPS TemCamXF416 digital camera (TVIPS GmbH, Gauting, Germany) using the SerialEM software package. Micrographs were taken as single tilt series over a range of −60° to +60° using SerialEM at tilt angle increment of 1°. Tomogram reconstruction was done with IMOD software.

### Gene expression analysis

To study gene expression plants were grown on half-strength MS plates covered with nylon mesh. Only root parts (around 100 mg) were collected, and total RNA was extracted using a Trizol-adapted ReliaPrep RNA Tissue Miniprep Kit (Promega). The DNase treatment was performed according to the manufacturer’s recommendations. Reverse transcription was carried out with oligo(dT) primers using Moloney murine leukemia virus reverse transcriptase according to the manufacturer’s instructions (Promega). The qPCR reaction was performed on an Applied Biosystems QuantStudio3 thermocycler using a SYBR™ Green PCR Master Mix (Thermo Fisher scientific), in a 96-well format. All transcripts were normalized to the housekeeping gene SAND (At2g28390) expression, using the qGene protocol (57). All primers used for qPCR are shown in Supplementary Table 1.

### Quantification and statistical analysis

For quantifying the Fluorol Yellow occupancy, confocal images were analyzed with the Fiji package (v.2.0.0-rc69/1.52p (build: 269a0ad53f); http://fiji.sc/Fiji (58). Contrast and brightness were adjusted in the same manner for all images. The suberized regions of the roots were measured together with total root lengths to determine the percentage of suberin occupancy. Relative suberization was defined by measurements of the suberin zone together with the circumference of the entire cell. Two-way ANOVA with Tukey HSD and Students t-test was subsequently used as a multiple-comparison procedure. Details about the statistical approaches used can be found in the Fig. legends. The data are presented as mean ± standard deviation (SD). All statistical analyses were done with the GraphPad Prism software v.9.0.0 (www.graphpad.com). Each experiment was repeated 2–4 times.

## Supporting information

all SFigures

Supplemental Movies description

SMovie 1

SMovie 2

SMovie 3

SMovie 4

SMovie 5

SMovie 6

SMovie 7

SMovie 8

SMethod info and primer list

## Acknowledgements

We thank Nasim Farahani Zayas for her contribution to generating the *proGPAT1:nls-GFP* line. Particular thanks go to Marie Barberon, Niko Geldner, and Robertas Ursache for discussions, sharing of unpublished information, and critical comments on the manuscript. Furthermore, we thank Philippe Reymond for helpful suggestions. The NASC stock center, Fred Beisson, and Niko Geldner are thanked for providing seed stocks and vectors. Furthermore, we thank Christel Genoud and Arnaud Paradis for leading the Electron Microscopy Facility and the Imaging Facility of UNIL, respectively. We are grateful to Sabine Freitag for expert technical assistance. This work was supported by the Swiss National Science Foundation (grant 31003A_170127 and 310030_188672 to CN) and by the Deutsche Forschungsgemeinschaft (DFG, INST 186/1167-1 to IF).

## Notes

### Competing Interest Statement

The authors have declared no competing interest.

