## Supplementary material for "The *GPAT4*/*6*/*8* clade functions in Arabidopsis root suberization non-redundantly with the *GPAT5/7* clade required for suberin lamellae": all SFigures

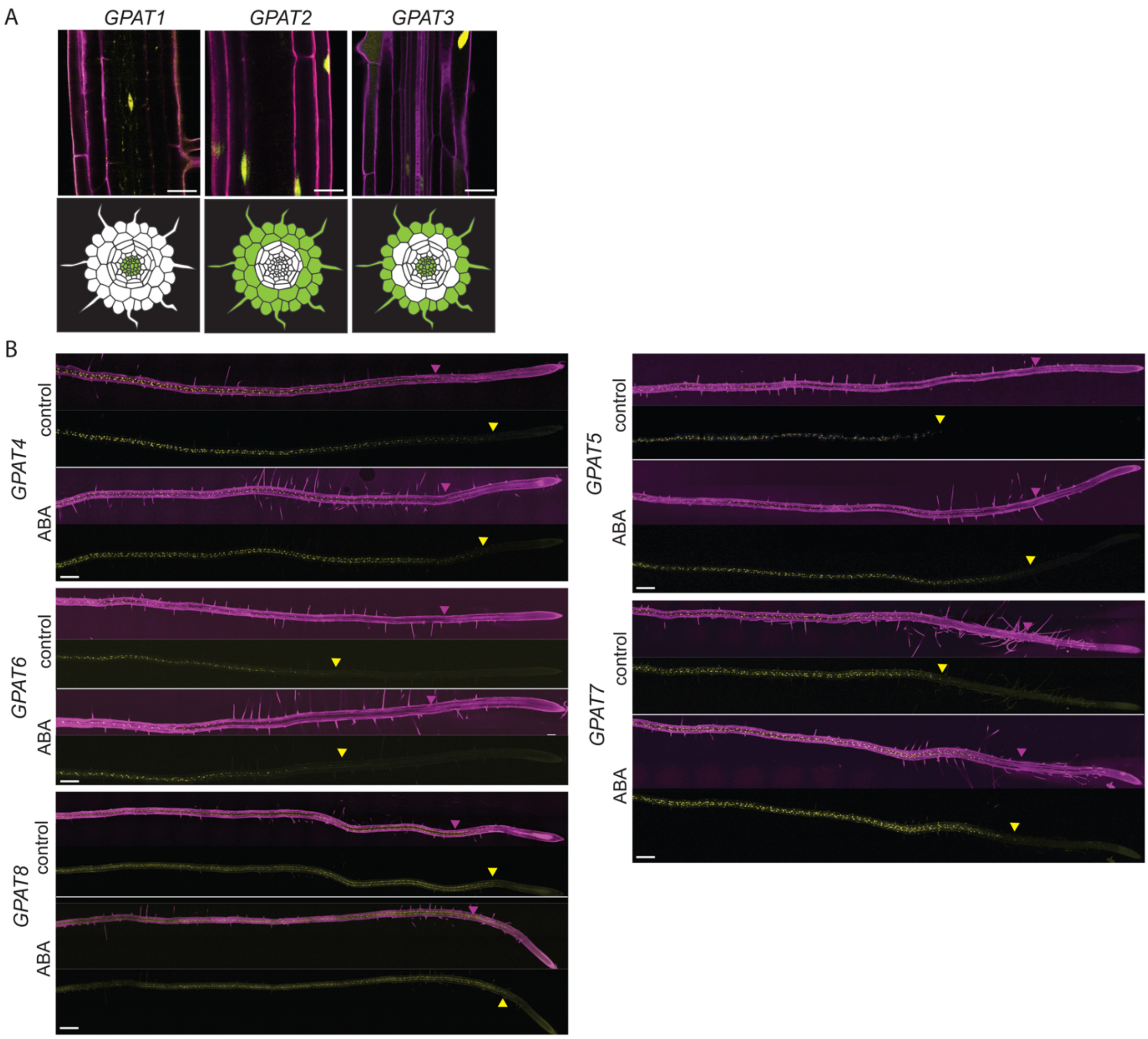


**Fig. S1. Cell-specific expression of *GPAT1-3* and ABA-regulated expression of *GPAT4-8***

(A) *PromoterGPATx::nls-GFP-GUS* reporter expression of *GPAT1*, *GPAT2* and *GPAT3* in transgenic Arabidopsis seedlings. GFP fluorescence is shown in yellow, propidium iodide (PI) fluorescence, as indicator of root cell shape, is presented in magenta. A fully suberized root section closest to the hypocotyl was evaluated in root of 5-day-old seedlings grown on half-strength MS medium. 3D-Z projections of longitudinal views (upper panels) and schematic diagrams of cross-section views are presented (lower panels). Scale bar 100 µm. (B) Abscisic acid (ABA)-regulation of *GPATs*. GFP-fluorescence (yellow) in transgenic plants expressing the indicated *promoterGPATx::nls-GFP-GUS* reporter gene fusion. Entire roots of 5-day-old seedlings that were grown for 4 days on half-strength MS medium and then transferred to half strength MS supplemented with 1 µM ABA or methanol (control) for 20 h were evaluated. 3D-Z projections of the florescence in the entire root are shown. Propidium iodide (PI) fluorescence (magenta) is presented as an indicator of cell layers and of the barrier properties of the cell walls. The yellow arrowhead indicates the position of the youngest cell showing GFP expression (in yellow). The magenta arrowhead indicates the position of the beginning of the blockage of PI into vasculature due to a fully developed Casparian strip. Scale bar 200 µm.

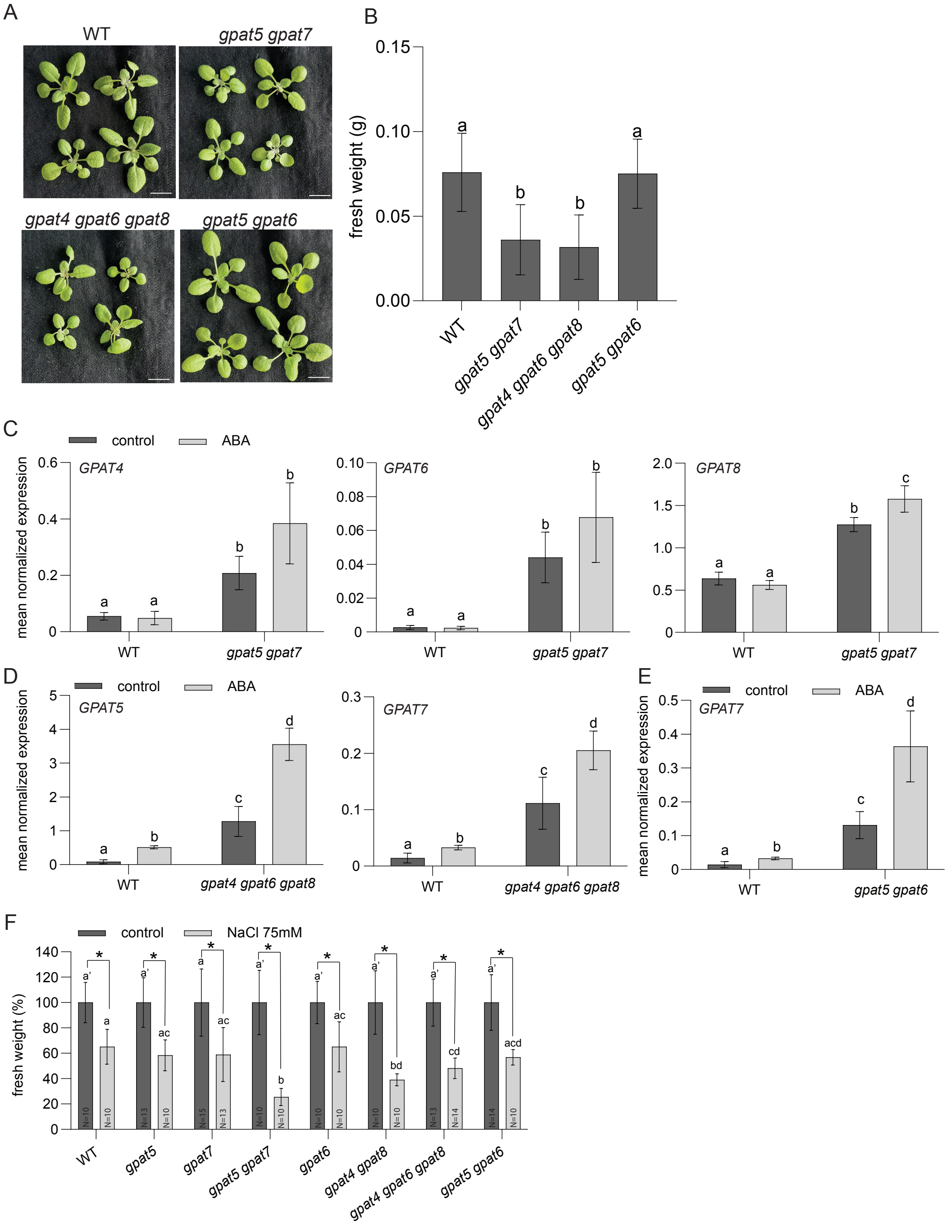


**Fig. S2. Growth habit and upregulation of functional *GPATs* in *GPAT* clade mutants**

(A-B) Growth habit of 3-week-old *GPAT* clade mutants and WT plants grown on soil. Macroscopic pictures of the rosettes (A) and their shoot fresh weight determination (B) revealed reduced growth in the pure clade mutant *gpat5 gpat7* and *gpat4 gpat6 gpat8*. Scale bars 1cm. Values represent the means +/- SD, n >10. Letters indicate significant differences, as determined by ANOVA with Tukey's post-hoc test, $p < 0.05$. (C-E) Expression of *GPAT*s in control conditions and after abscisic acid (ABA) treatment in *gpat5 gpat7* (C), *gpat4 gpat6 gpat8* (D), and *gpat5 gpat6* (E). *GPAT* expression was evaluated by RT-qPCR in 5-day-old roots that were grown for 4 days on half-strength MS medium and then transferred to half-strength MS medium supplemented with 1 μM ABA or methanol (control) for 20h. Values represent means ± SD, n = 3. Different lowercase letters indicate significant differences relative to the WT, as determined by ANOVA and Tukey's post-hoc test, $p < 0.05$. (F) Fresh weight reduction in *gpat* mutants under salt stress as shown in Fig1. Plants were grown for 4 days on half-strength MS medium and then transferred to half-strength MS medium containing 75 mM NaCl for 9 days. Mean of fresh weight ± SD, n ≥ 10. Different lowercase letters indicate significant differences, as determined by ANOVA and Tukey's post hoc test, *p*-value < 0.05. Significant differences between salt stress and control conditions are shown by an asterisk, as determined by Student's t-test *: $p < 0.05$.

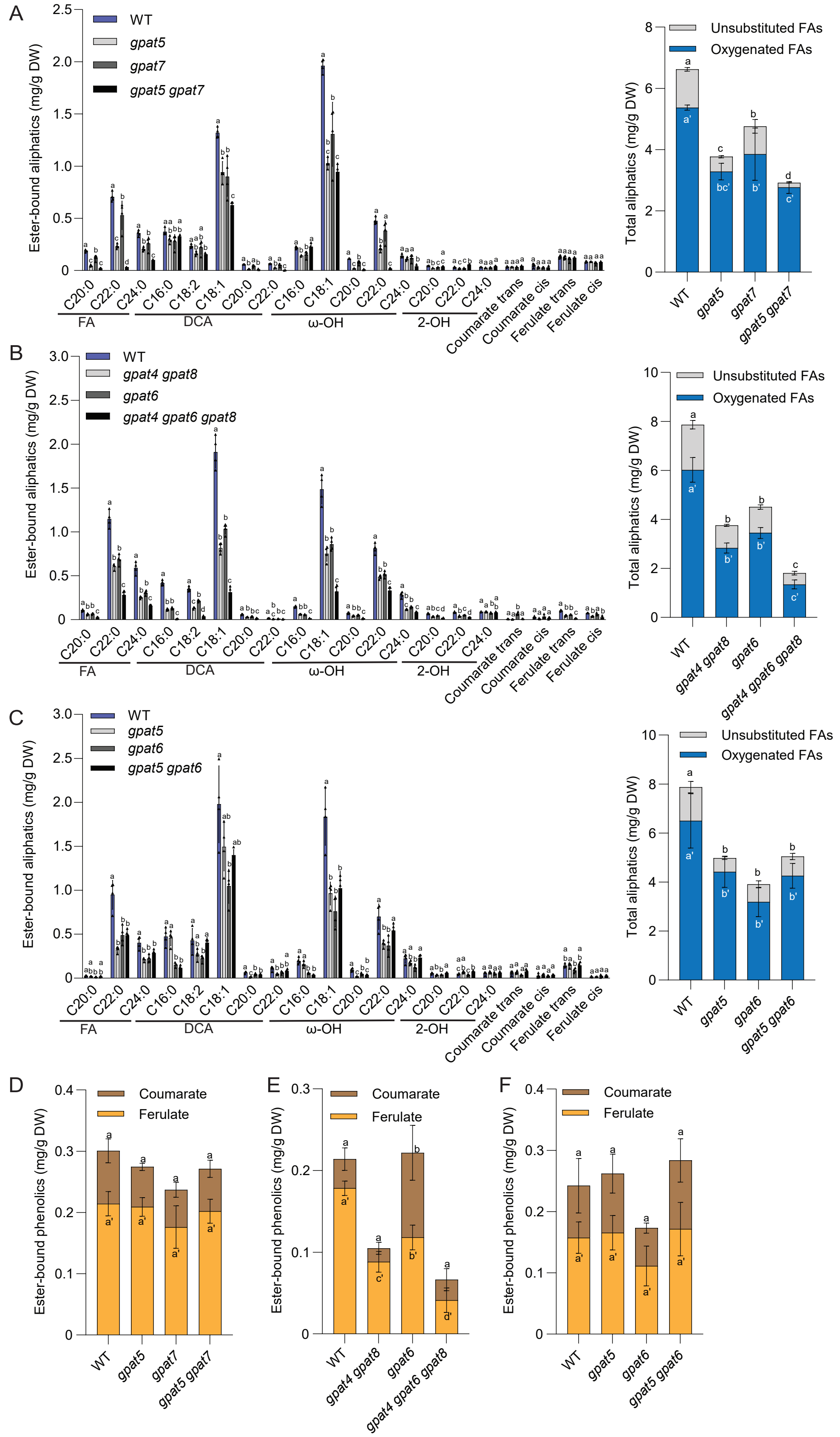


**Fig. S3. Characteristic alterations of endodermal suberin monomer composition in *GPAT clade* mutants**

(A-C) Quantification of aliphatic and aromatic ester-bond suberin monomers of *GPAT5/7*-clade mutants (A), *GPAT4/6/8*-clade mutants (B) and mixed clade mutants (C) with their respective wild type (WT) control. All identified aliphatic and aromatic suberin monomers are presented on the left with the total of identified aliphatic compounds grouped by substance classes on the right. (D-F) The total of the principal ester-bound hydroxycinnamic acids of *GPAT5/7* clade mutants (D), *GPAT4/6/8*-clade mutants (E) and mixed clade mutants (F). Roots of 5-day-old plants grown on half-strength MS medium were analyzed. Values represent the means ± SD, n = 4. Different lowercase letters indicate significant differences among the genotypes, as determined by ANOVA with Tukey's post hoc test, $p$-value < 0.05; Black triangles in the graphs on the left (A-C) depict individual data points.  FA, fatty acid; DCA; dicarboxylic acid; DW, dry weight.

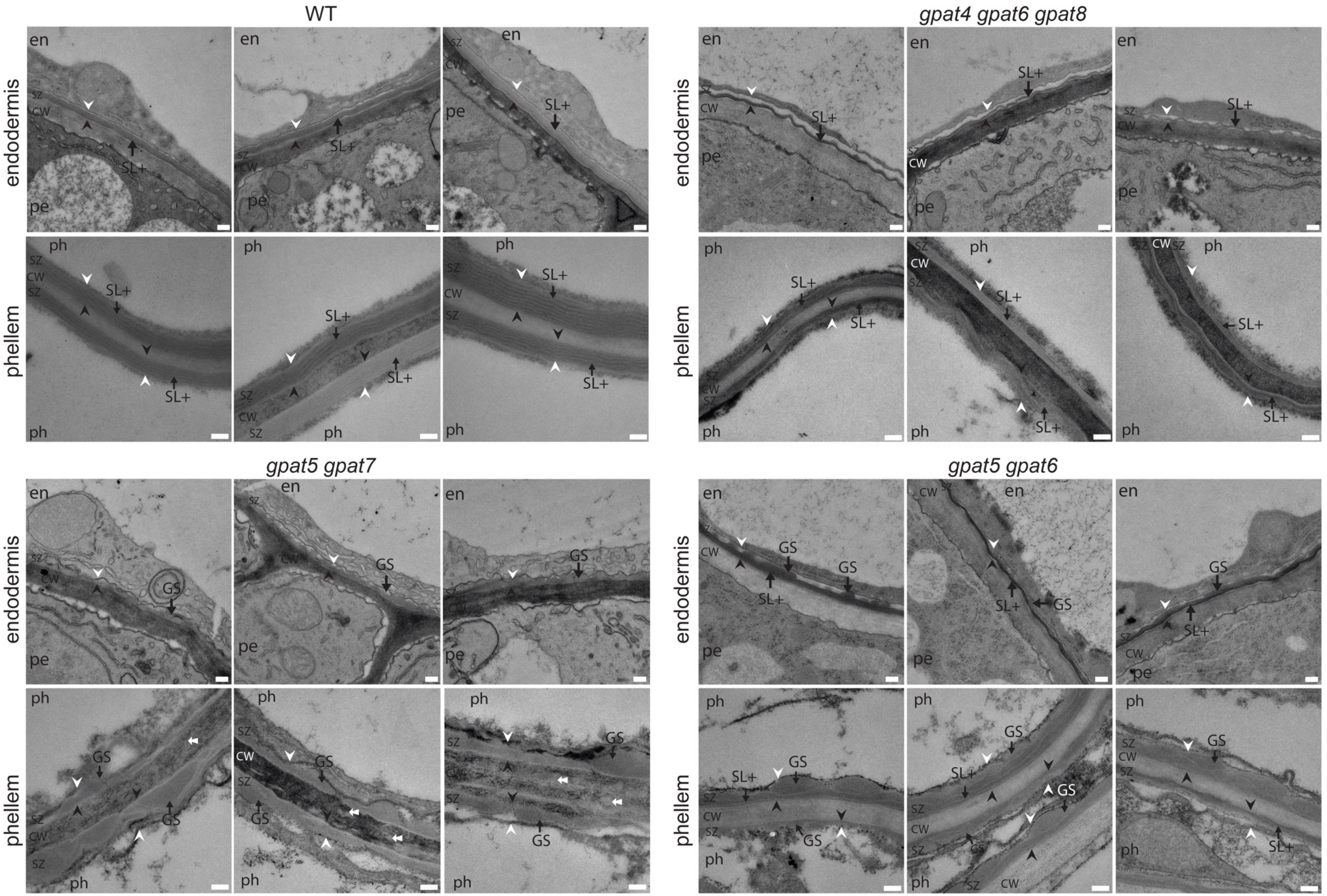
WT
gpat4 gpat6 gpat8
gpat5 gpat7
gpat5 gpat6
endodermis
phellem
en
pe
ph
sz
cw
SL+
GS

**Fig. S4. Characteristic alterations of the suberin ultrastructure in *GPAT*-clade mutants.**

Suberin ultrastructure was analyzed in endodermal cells and phellem cells by TEM. Pictures show three areas of different roots of WT and *gpat* mutants, in addition to these shown in Fig. 3. Scale bars represent 200 µm for pictures of the endodermal cells and 100 µm of phellem cells. CW, cell wall; SZ, suberin zone; white arrowhead, plasma membrane; black arrowhead, junction between suberin zone (SZ) and polysaccharide cell wall (CW). Black arrows point to polyester depositions of different structure: GS, globular suberin; SL+, suberin layer strongly lamellated substructure; white arrows point to globular suberin that is embedded in the polysaccharide cell wall; en, endodermis cell; pe, pericycle cell; ph, phellem cell.

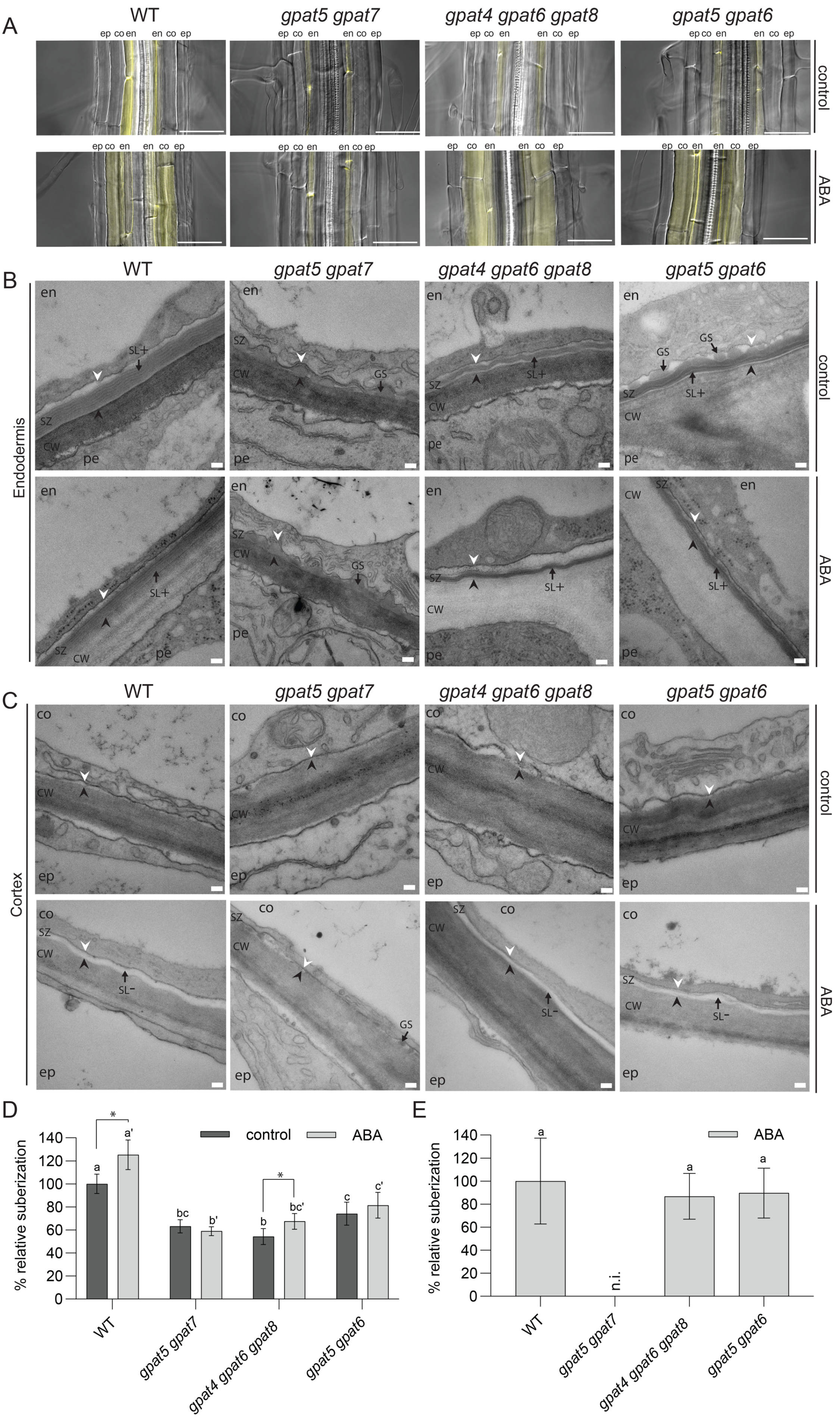
A
WT
gpat5 gpat7
gpat4 gpat6 gpat8
gpat5 gpat6
ep co en en co ep
control
ABA
B
Endodermis
en
pe
SZ
CW
SL+
GS
C
Cortex
co
ep
SL-
D
% relative suberization
control
ABA
*
a
a'
bc
b'
b
bc'
c
c'
0
20
40
60
80
100
120
140
E
n.i.

**Fig. S5. Suberization in *GPAT* clade mutants after ABA treatment**

(A-C) Cell type-specific suberization and suberin ultrastructure was characterized in different cell types in 5-day-old seedlings. (A) Fluorol Yellow staining showed different degrees of root suberization under control conditions (upper row) and after ABA treatment (lower row) in *GPAT* clade mutants in comparison to the wild type (WT). (B) The ultrastructure of suberin was assessed by transmission electron microscopy (TEM) in the endodermis on the side neighboring the pericycle under control conditions (upper row, pictures also shown in Fig. 3) and after ABA treatment (lower row) revealing specific modifications in each *gpat* mutant compared to the wild type (WT). (C) The ultrastructure of suberin was assessed in the cortex, on the side neighboring the epidermis, under control conditions (upper row) and after ABA treatment (lower row) revealing the lack of a suberizing layer after ABA treatment in *gpat5 gpat7*. Scale bars, 50 µm in pictures showing FY staining, and 100 nm in TEM pictures. CW, cell wall; SZ, suberin zone. White arrowhead, plasma membrane; black arrowhead, junction between suberin zone (SZ) and polysaccharide cell wall (CW). Black arrows point to polyester deposits of different structures: GS, globular suberin; SL+, suberin layer with a strongly lamellated substructure; SL-, largely amorphous suberin layer with little substructure; en, endodermal cell; ep, epidermal cell, pe, pericycle cell; co, cortical cell. (D-E) Quantification of different ultrastructural features visualized by TEM. The thickness of the suberized zone seen in the endodermis (D) or in cortical cells (E) was quantified by measuring the area between the PM and CW in six cells. Values are represented relative to the WT grown in control conditions (%). A continuous suberized layer could not be identified (n. i.) in cortical cells of *gpat5 gpat7* after ABA treatment. Different lowercase letters indicate significant differences between the suberized areas of different genotypes, as determined by ANOVA with Tukey's post-hoc test, $p < 0.05$ and $n = 6$. Significant differences between the control and ABA treatments for each genotype are indicated by an asterisk (*), as determined by Student's t-test, $p < 0.05$.

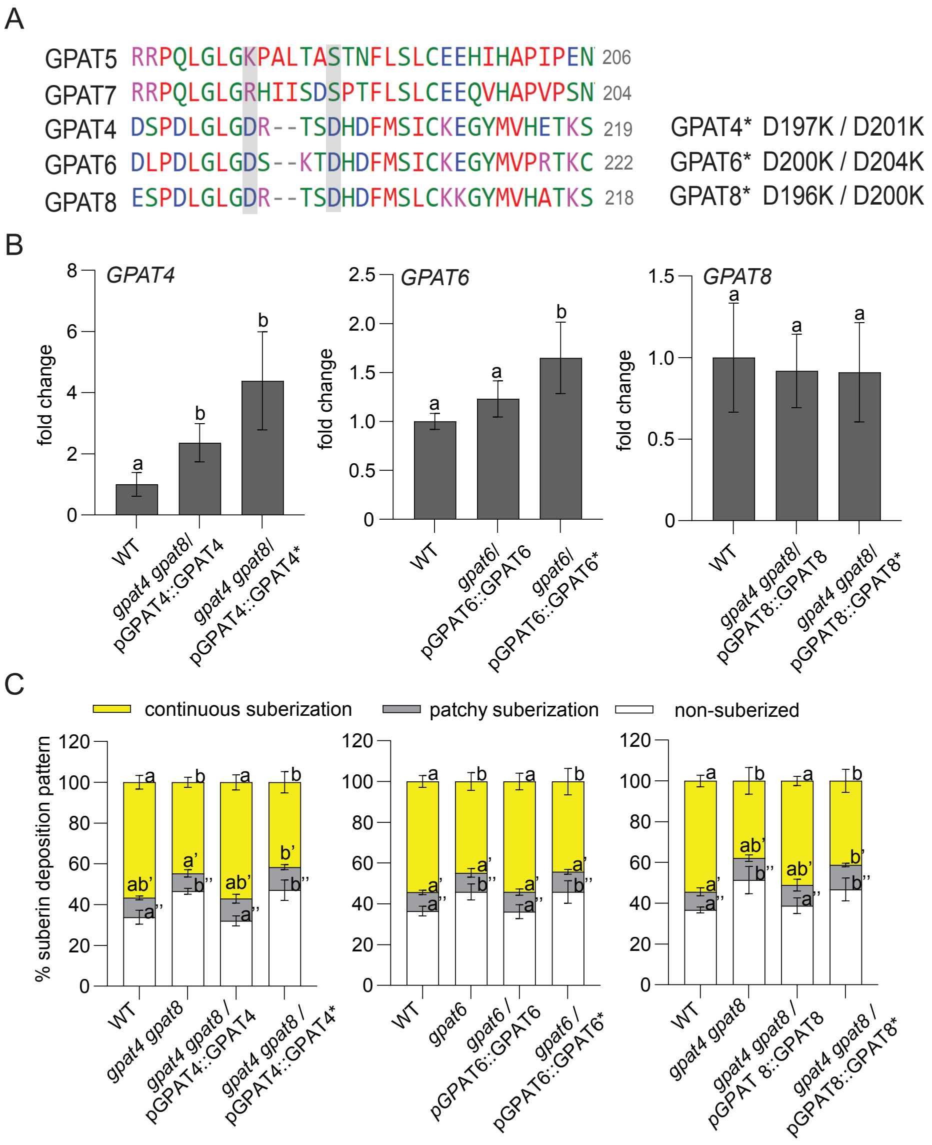


**Fig. S6. Effect of mutations in the phosphatase domain of GPAT4, GPAT6, and GPAT8 on suberization**

(A) Schematic presentation of mutations introduced in motive III of the phosphatase of GPAT4/6/8. Two conserved aspartic acid (D) residues of the phosphatase domain in GPAT4, GPAT6, and GPAT8 were mutated to lysine (K) residues (GPAT4*, GPAT6*, and GPAT8*). Numbers in gray indicate the position of motive III in the amino acid sequence of the respective protein. (B) Expression of mutated and unmutated *GPAT* genes in comparison to the respective wild type (WT) control. Transgenic lines were selected that had a similar or even higher expression of the respective *GPAT* gene than in the wild type (WT). Expression of *GPATs* was evaluated by RT-qPCR in roots of 5-day-old plants grown on half-strength MS medium. Results are presented as fold-changes in comparison to the WT based on three biological replicates. Values represent the means ± SD, n=3. Different lowercase letters indicate significant differences, as determined by ANOVA with Tukey's post-hoc test, $p < 0.05$. (C) Suberin coverage was not complemented by *GPATs* having a mutated phosphatase domain. Fluorol Yellow (FY) staining was evaluated in the wild type (WT), *gpat* mutants and *gpat* mutants transformed either with the respective unmutated or mutated *GPAT* gene. Seedlings were grown for 5 days on half-strength MS medium under standard conditions. Quantification of the suberin pattern along the longitudinal axis of the root was performed. Data are means ± SD, n ≥ 6. Different letters indicate significant differences, as determined by ANOVA with Tukey's post-hoc test, $p < 0.05$.

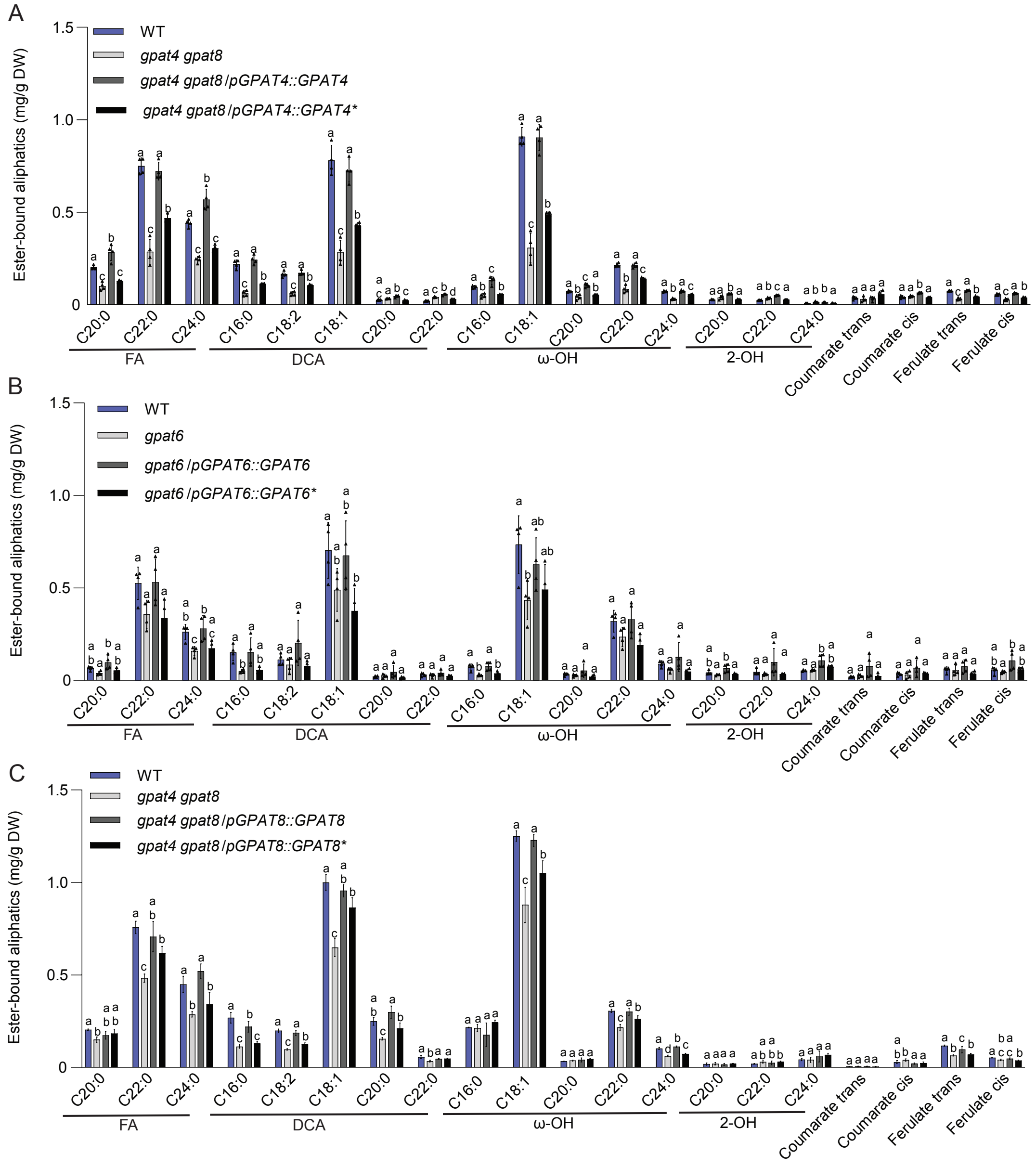


**Fig. S7. Reduced suberin amount by phosphatase domain-mutated GPATs**

(A-C) Effect of mutating the phosphatase domain of GPATs, i.e. GPAT4* (A) GPAT6* (B) and GPAT8* (C) on suberin formation. Graphs show quantification of aliphatic and aromatic ester-bond suberin monomers isolated from 5-day-old seedlings.of *gpat* mutants as well as transgenic *gpat* mutant lines complemented with the respective unmutated *GPAT* gene or phosphatase domain mutated *GPAT* gene (*GPAT**) in comparison to the wild type (WT). Values represent the means ± SD, n = 4. Different lowercase letters indicate significant differences between genotypes, as determined by ANOVA with Tukey's post hoc test, $p < 0.05$; Black triangles depict individual data points. FA, fatty acid; DCA; dicarboxylic acid; DW, dry weight.

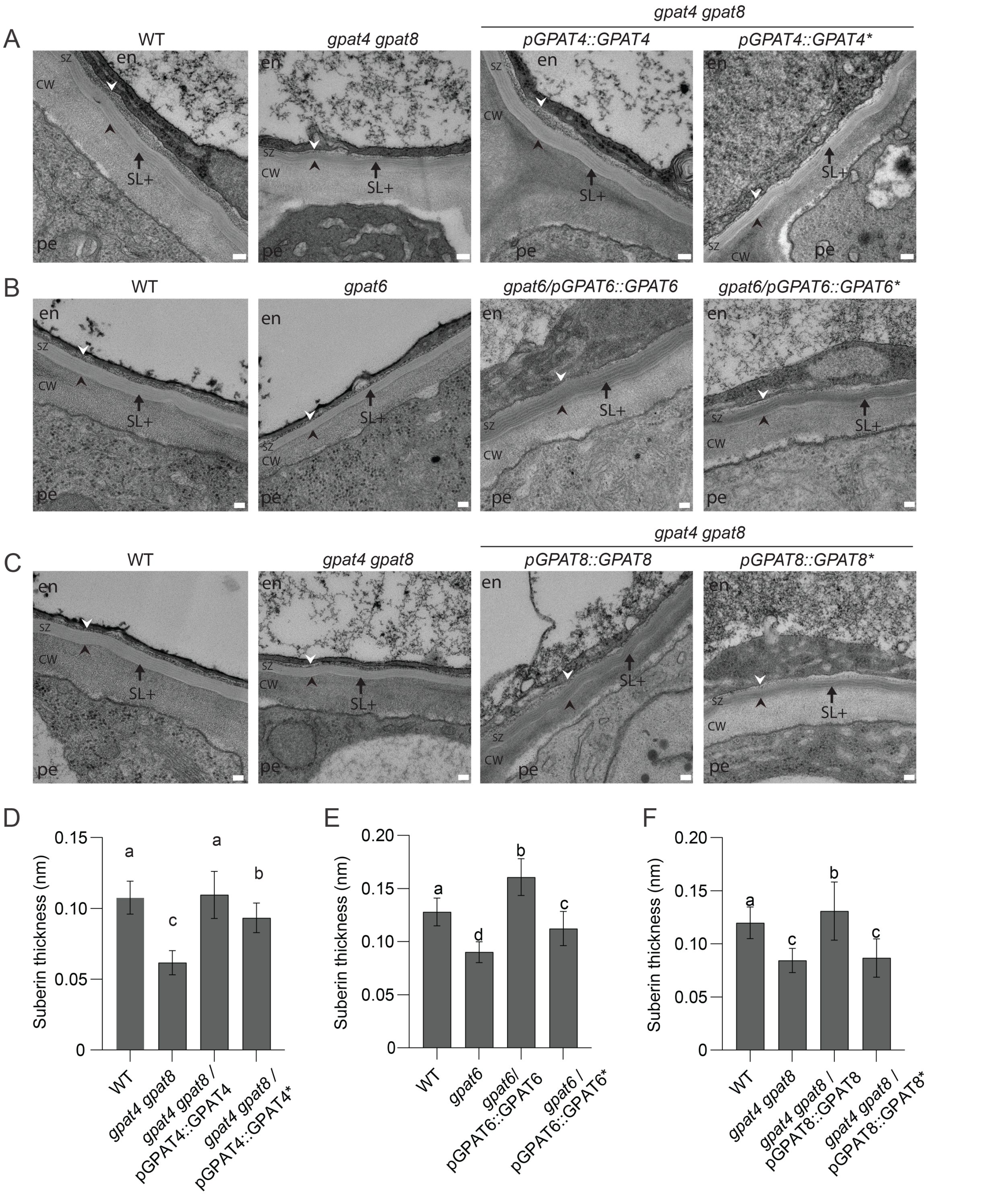


**Fig. S8. Effect of mutations in the phosphatase domain of GPAT4, GPAT6 and GPAT8 on suberin ultrastructure**

(A-C) Lamellated suberin ultrastructure in the endodermis on the side neighboring the pericycle in wild type (WT), *gpat4 gpat8* as well as *gpat4 gpat8* transformed with the unmutated *GPAT4* gene or phosphatase domain mutated *GPAT4* gene (*GPAT4**) (A) *gpat6* with the unmutated *GPAT6* gene or phosphatase domain mutated *GPAT6* gene (*GPAT6**) (B) and *gpat4 gpat8* transformed with the unmutated *GPAT8* gene or phosphatase domain mutated *GPAT8* gene (*GPAT8**) (C). Scale bar 100 nm. CW, cell wall; SZ, suberin zone. White arrowhead, plasma membrane; black arrowhead, junction between suberin zone (SZ) and polysaccharide cell wall (CW). Black arrows point to polyester deposits: SL+, suberin layer with a lamellated substructure. (D-F) The phosphatase mutated *GPATs* only partially complemented the thickness of the suberin layer. The thickness of the suberin layer visualized in TEM was measured in at least 4 endodermal cells by measuring the thickness of the suberin lamellae. Values represent the means ± SD, $n \geq 50$. Different lowercase letters indicate significant differences among the genotypes, as determined by ANOVA with Tukey's post hoc test, $p$-value $< 0.05$.

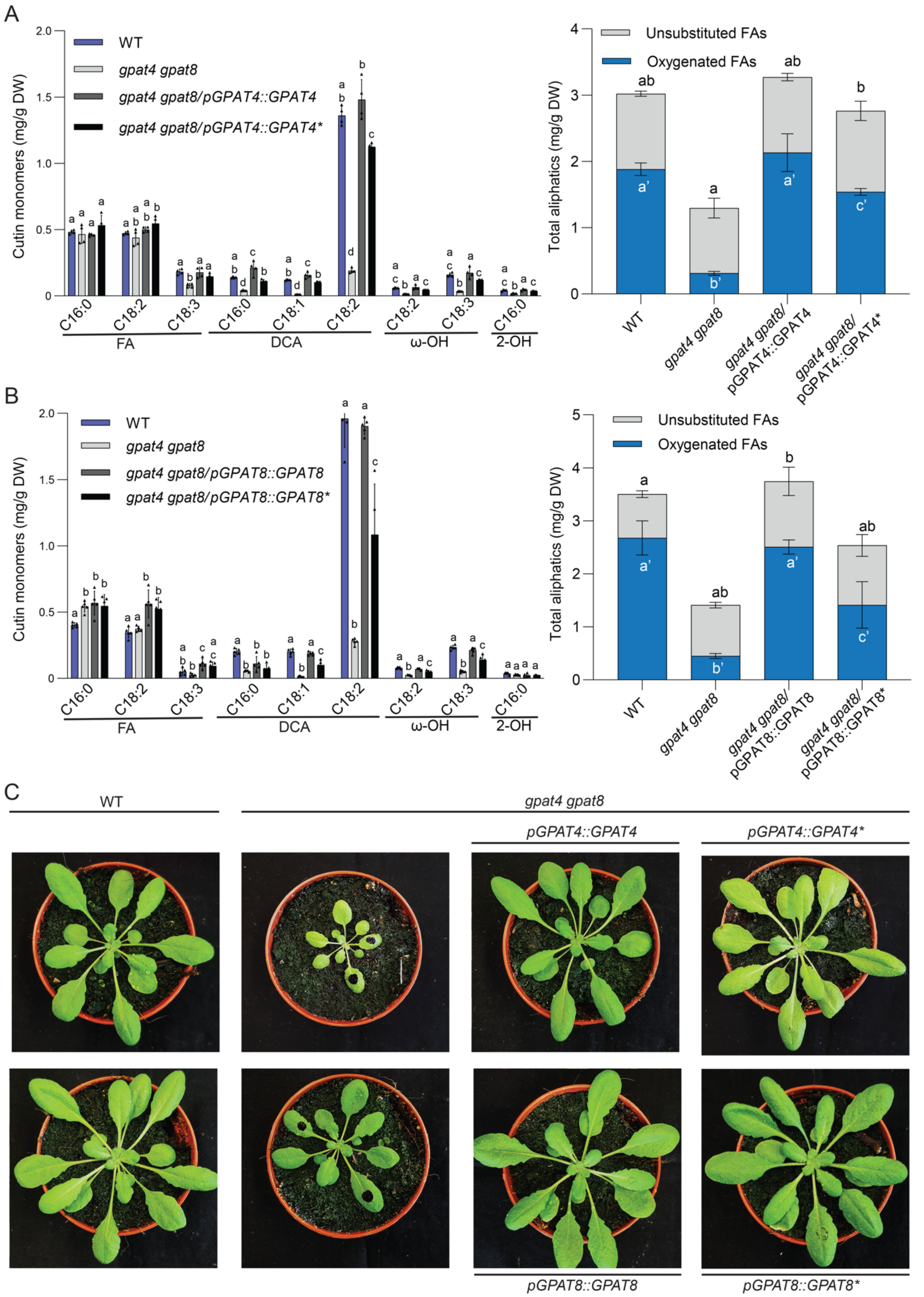


**Fig. S9. Partial complementation of cutin amounts in *gpat4 gpat8* by phosphatase domain-mutated GPAT4 and GPAT8**

(A-B) Effect of mutating the phosphatase domain of GPATs on cutin formation. In contrast to *GPAT4*, *GPAT4** complements only partially the deficiencies in cutin biosynthesis in rosette leaves of the *gpat4 gpat8 double* mutant (A). In contrast to *GPAT8*, *GPAT8** complements only partially the deficiencies in cutin biosynthesis in the *gpat4 gpat8* mutant (B). The composition of the cutin monomers are shown on the left (A, B) and the total of analyzed cutin monomers, grouped by substance classes are shown on the right (A, B). Entire rosettes of 4 weeks-old soil-grown plants were harvested and analyzed by GC-MS. Values represent the means ± SD, n = 4. Different lowercase letters indicate significant differences between genotypes, as determined by ANOVA with Tukey‘s post hoc test, $p < 0.05$. Black triangles in the graphs on the left depict individual data points. FA, fatty acid; DCA; dicarboxylic acid; DW, dry weight. (C) Evaluation of the permeability of the cuticle in rosette leaves by Toluidine Blue (TB) staining. Droplets of 10 µl TB were incubated for 1 h on two leaves (leaf 5-6 and leaf 8-9) of 6 plants that were kept under high humidity and then removed by washing with water. TB staining of representative rosettes were photographed showing that the barrier properties were largely restored by GPAT4* and GPAT8*, respectively, despite the deficiencies in cutin amount.

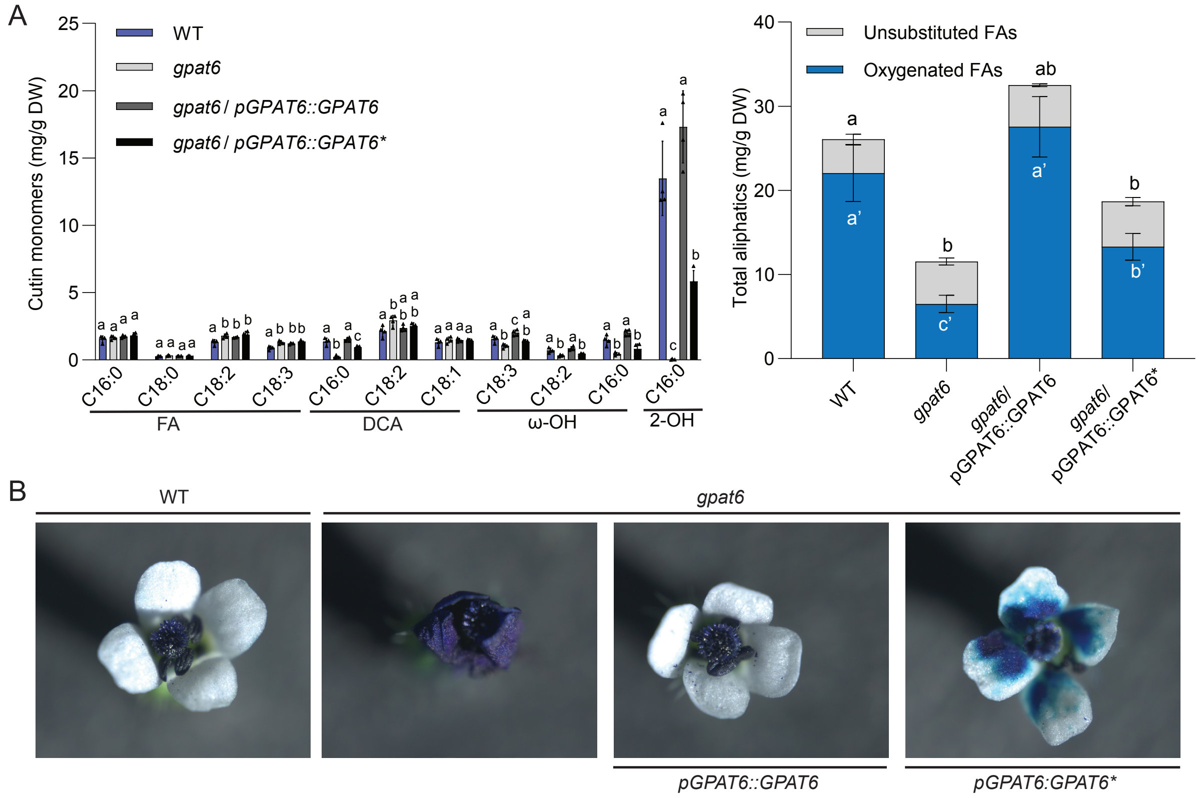


**Fig. S10. Partial complementation of cuticle-related phenotypes in the *gpat6* mutant by phosphatase domain-mutated GPAT6**

(A) Effect of mutating the phosphatase domain of GPAT6 on cutin formation. In contrast to *GPAT6*, *GPAT6** complements only partially the deficiencies in cutin biosynthesis in *gpat6* mutant flowers. The composition of the cutin monomers is shown on the left and the total of analyzed cutin monomers grouped by substance classes are shown on the right. 20 flowers having 4 petals were harvested and analyzed by GC-MS. Values represent the means ± SD, n = 4. Different lowercase letters indicate significant differences to wild type (WT), as determined by ANOVA with Tukey's post hoc test, $p < 0.05$; Black triangles in the graph on the left depict individual data points. FA, fatty acid; DCA, dicarboxylic acid; DW, dry weight. (B) Evaluation of the permeability of the petal cuticle by Toluidine Blue (TB) staining. 20 flowers were incubated in TB for 90 min and then washed in water. Representative flowers were photographed showing that the barrier properties were only partially restored by GPAT6*.

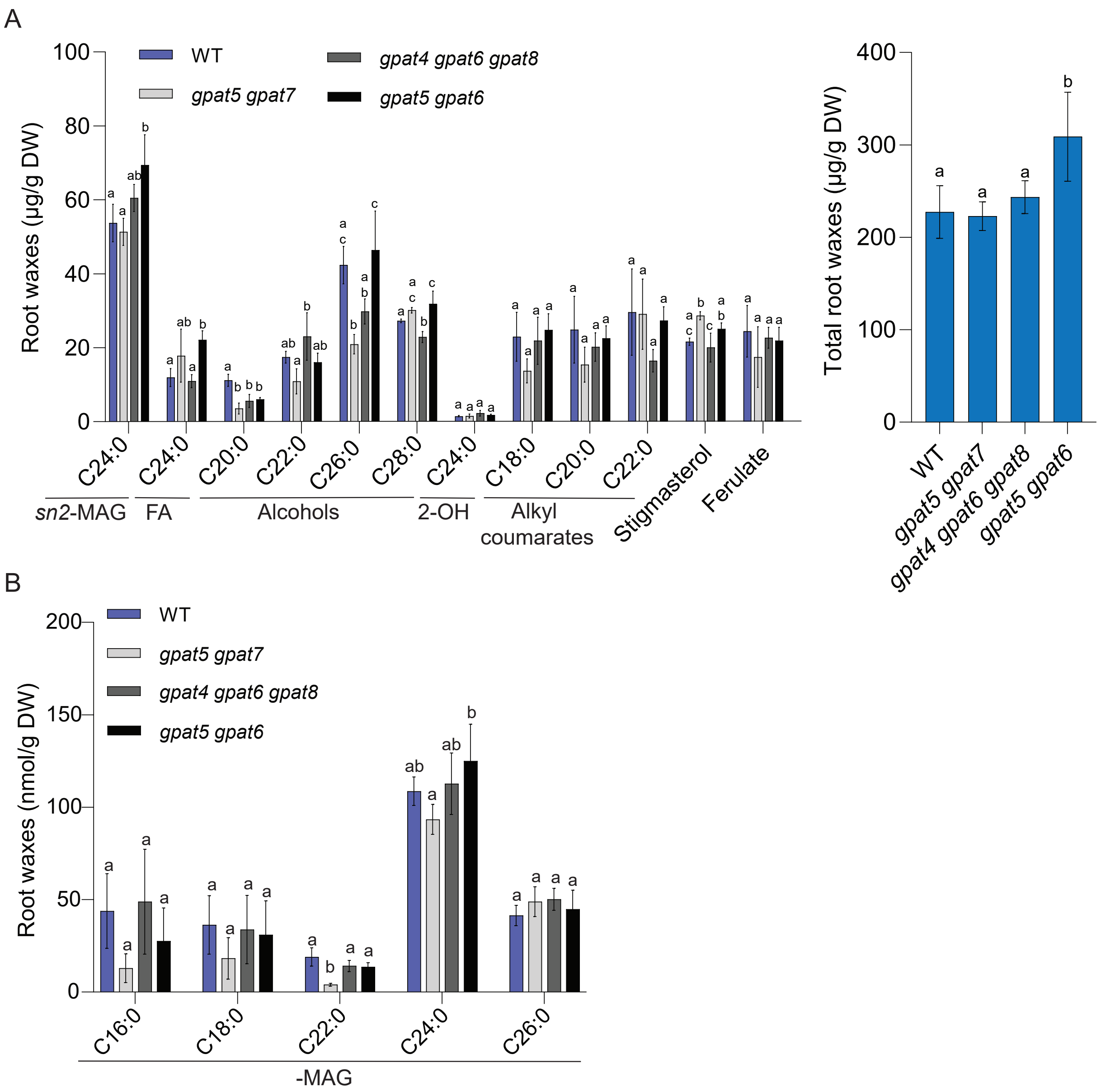


**Fig. S11. Characterization of root wax composition in *GPAT* clade mutants**

(A-B) Quantification of root waxes isolated from roots of 3-week-old *GPAT* clade mutants in comparison to the wild type (WT). (A) All root-wax compounds, identified by GC-MS/FID on the left are presented as well as the sum of the evaluated root waxes on the right. (B) Detailed evaluation of monoacylglycerols from the same preparation as shown in A, identified by targeted LC-MS/MS approach. Plants were grown on half-strength MS medium. Values represent means ± SD, n = 4. Different lowercase letters indicate significant differences, as determined by ANOVA with Tukey's post-hoc test, $p < 0.05$. FA, fatty acid; MAG; monoacylglycerol; DW, dry weight.

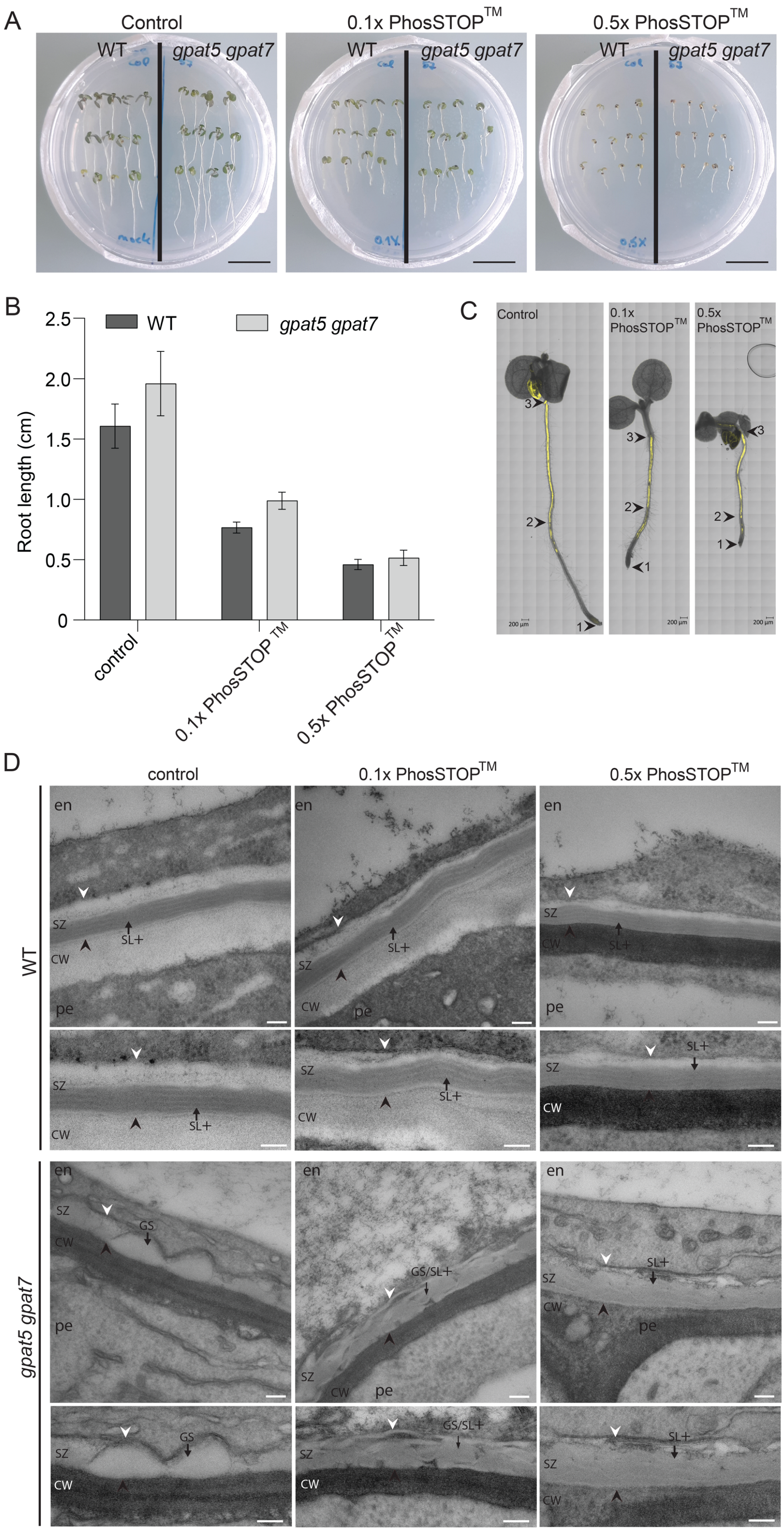

A
Control
0.1x PhosSTOP™
0.5x PhosSTOP™
WT
gpat5 gpat7
B
WT
gpat5 gpat7
Root length (cm)
2.5
2.0
1.5
1.0
0.5
0
control
0.1x PhosSTOP™
0.5x PhosSTOP™
C
Control
0.1x PhosSTOP™
0.5x PhosSTOP™
D
control
0.1x PhosSTOP™
0.5x PhosSTOP™
WT
gpat5 gpat7
en
pe
SZ
CW
SL+
GS
GS/SL+

**Fig. S12. Effect of PhosSTOP™ treatment on plant growth and suberization**

(A-C) Assessment of growth and suberization of seedlings after application of the phosphatase inhibitor PhosSTOP™ (A) Pictures of wild type (WT) and *gpat5 gpat7* seedlings grown for three days on half-strength MS medium and for 2 days on half-strength MS supplemented with indicated concentrations of the phosphatase inhibitor PhosSTOP™. Scale bars represent 1 cm. (B) Root length of wild type (WT) and *gpat5 gpat7* mutant seedlings grown as described in A. No statistical difference was observed between the mutant and the WT under each growth condition. (C) Suberin, as evaluated by Fluorol Yellow (FY) staining, was deposited in wild type (WT) seedlings after growth in the presence of the indicated PhoSTOP™ concentrations (as shown in A). Arrowhead 1 indicates location of the root tip, arrowhead 2 the start of continuous suberization and arrowhead 3 the location of the root/ hypocotyl junction. Scale bars represent 200 µm. (D) The ultrastructure of suberin was assessed by transmission electron microscopy (TEM) in the endodermis of wildtype (upper panel) as well as *gpat5 gpat7* (lower panel) on the side neighboring the pericycle under control conditions and after phosphatase inhibitor PhosSTOP™ (0.1x and 0.5x) treatment (related to Fig. 4B; additional examples in Fig. S13). Higher magnifications are shown below each picture. CW, cell wall; SZ, suberin zone. White arrowhead, plasma membrane; black arrowhead, junction between suberin zone (SZ) and polysaccharide cell wall (CW). Arrows point to polyester deposits of different structures; GS: globular suberin; SL+: suberin layer with a strongly lamellated substructure; GS/SL+: intermediate structure with globular suberin as well as lamellated substructure. Scale bar 100 µm.

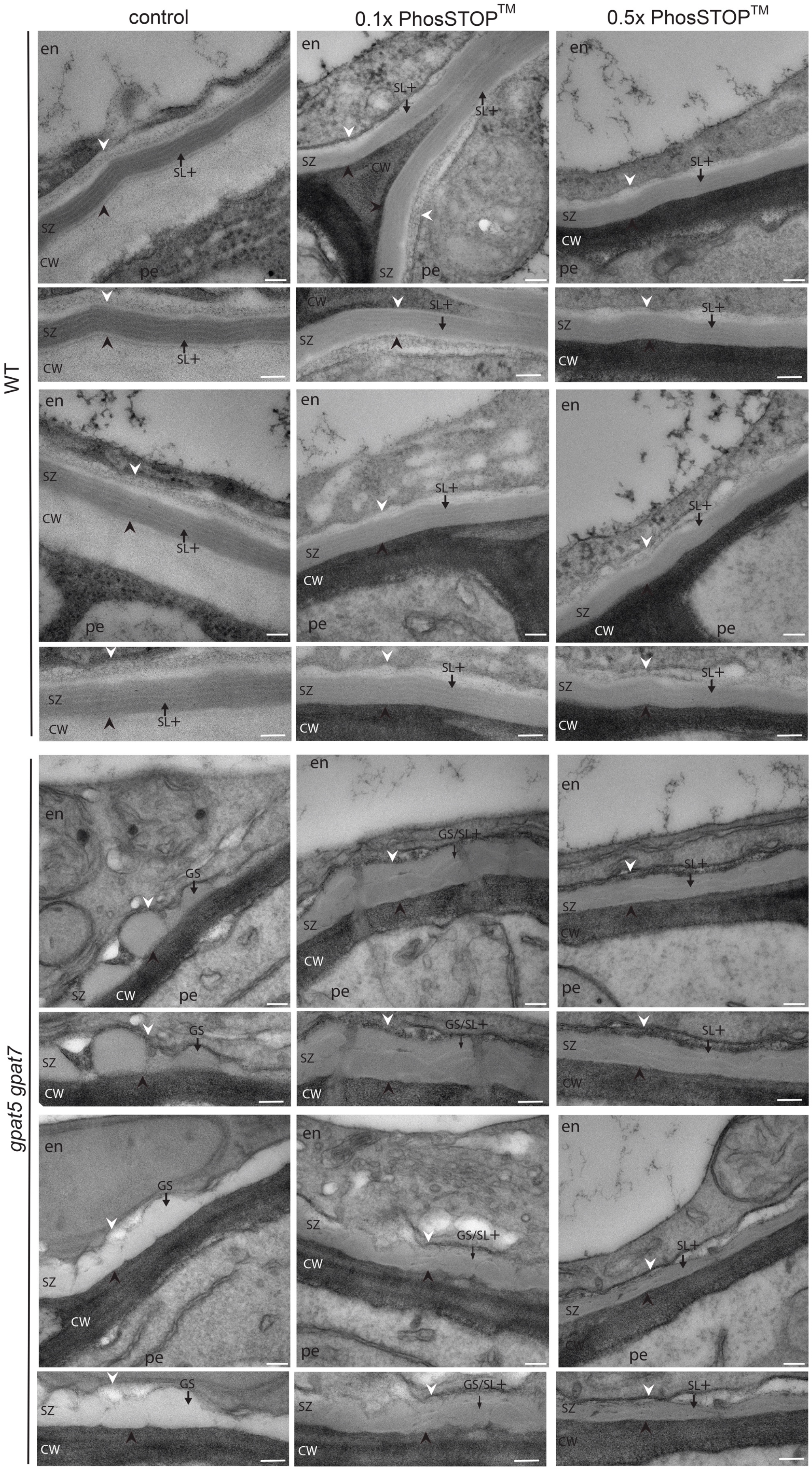
control
0.1x PhosSTOP™
0.5x PhosSTOP™
WT
gpat5 gpat7
en
SZ
CW
pe
SL+
GS
GS/SL+

**Fig. S13. Effect of PhosSTOP$^{TM}$ treatment on suberin ultrastructure**

The ultrastructure of suberin was assessed by transmission electron microscopy (TEM) in the endodermis of wildtype (upper panel) as well as *gpat5 gpat7* (lower panel) on the side neighboring the pericycle under control conditions and after phosphatase inhibitor PhosSTOP$^{TM}$ (0.1x and 0.5x) treatment. Two additional pictures are given of each genotype and treatment in addition to these shown in Fig. S12. Higher magnifications are shown below each picture. CW, cell wall; SZ, suberin zone. White arrowhead, plasma membrane; black arrowhead, junction between suberin zone (SZ) and polysaccharide cell wall (CW). Arrows point to polyester deposits of different structures; GS: globular suberin; SL+: suberin layer with a strongly lamellated substructure; GS/SL+: intermediate structure with globular suberin as well as lamellated substructure. Scale bar 100 µm.

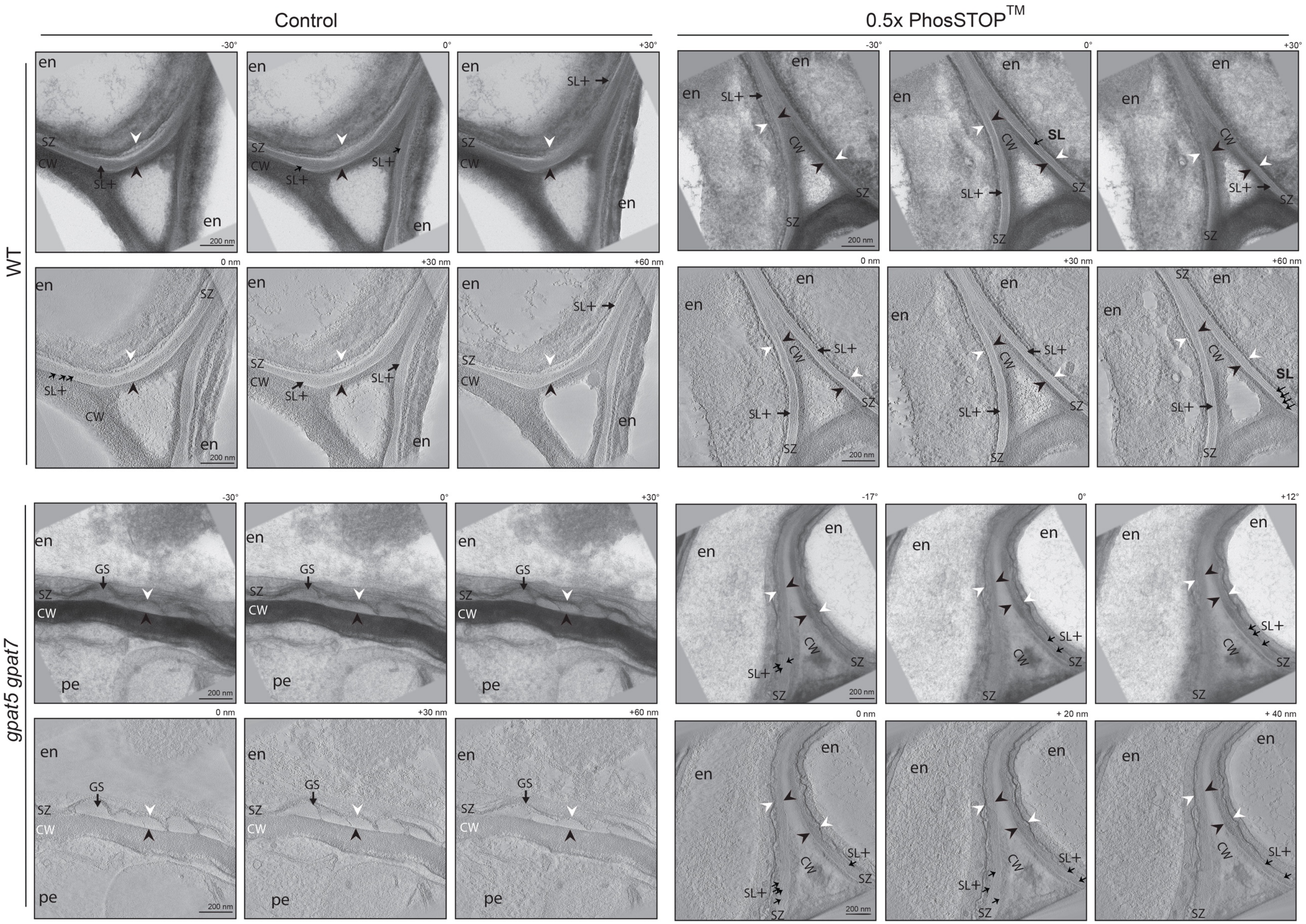
Control
0.5x PhosSTOP™
WT
gpat5 gpat7
-30°
0°
+30°
0 nm
+30 nm
+60 nm
-17°
+12°
+ 20 nm
+ 40 nm
en
pe
SZ
CW
SL+
SL
GS
200 nm

**Fig. S14. Effect of PhosSTOP$^{TM}$ treatment on the internal organization of suberin**

The internal organization of suberin was assessed by electron tomography in the endodermis of wildtype (upper panel) and the *gpat5 gpat7* double mutant (lower panel) on the side neighboring the pericycle under control conditions and after phosphatase inhibitor PhosSTOP$^{TM}$ (0.5x) treatment. Roots cut at 1 mm from hypocotyl were evaluated. The upper rows show three different tilt angles, respectively -30°, 0° and +30° from the tilt series (-60°, +60°). The lower rows show a series of three optical sections generated from the tomogram. Arrows point to polyester deposits of different structures; GS: globular suberin; SL+: suberin layer with a strongly lamellated substructure. White arrowhead, plasma membrane; black arrowhead, junction between suberin zone (SZ) and polysaccharide cell wall (CW). en, endodermis; pe, pericycle; CW, cell wall; SZ, suberin zone. Scale bars represent 200 nm.

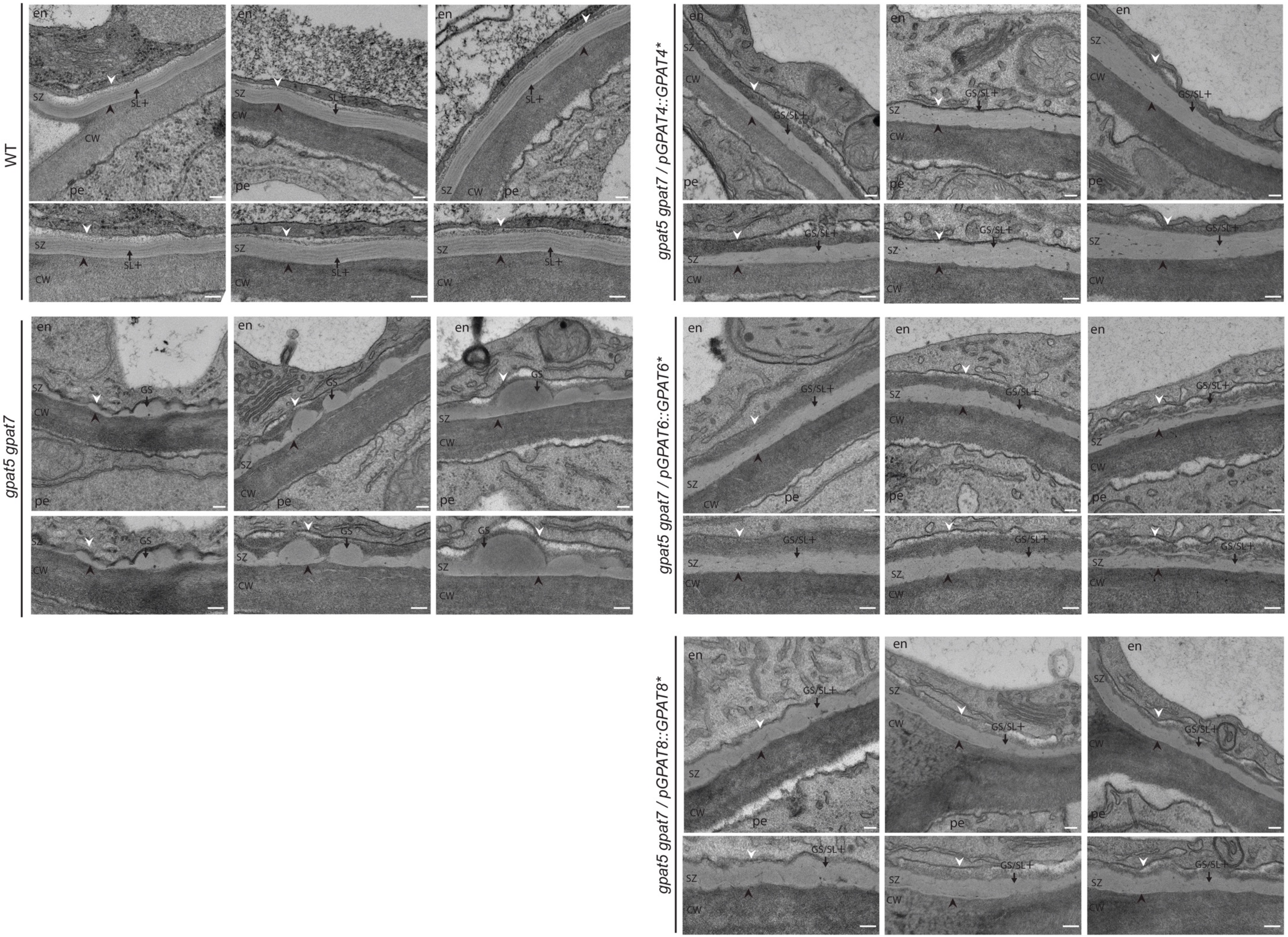
WT
gpat5 gpat7
gpat5 gpat7 / pGPAT4::GPAT4*
gpat5 gpat7 / pGPAT6::GPAT6*
gpat5 gpat7 / pGPAT8::GPAT8*
en
SZ
CW
pe
SL+
GS
GS/SL+

**Fig. S15. Effect of the expression of phosphatase-dead *GPAT4/6/8* clade members on suberin structure in the *GPAT5/7* clade mutant**

Transgenic *gpat5 gpat7* plants of the T1 generation expressing a mutated *GPAT* gene encoding for a member of the *GPAT4/6/8* clade, in which the two conserved aspartic acid residues of the phosphatase domain were mutated to lysine residues and expressed under their respective native promoter (pGPAT4::GPAT4*, pGPAT6::GPAT6*, pGPAT8::GPAT8*) as well as the wild type (WT) and *gpat5 gpat7* double mutant were evaluated. Different genotypes were grown for 5 days on half-strength medium. Representative pictures are shown here together with an enlargement underneath to highlight the suberin ultrastructure (see also Fig. 4B). CW, cell wall; SZ, suberin zone. White arrowhead, plasma membrane; black arrowhead, junction between suberin zone (SZ) and polysaccharide cell wall (CW). Arrows point to polyester deposits of different structures; GS: globular suberin; SL+: suberin layer with a strongly lamellated substructure; GS/SL+: intermediate structure with globular suberin as well as lamellated substructure. Scale bar, 100 µm.

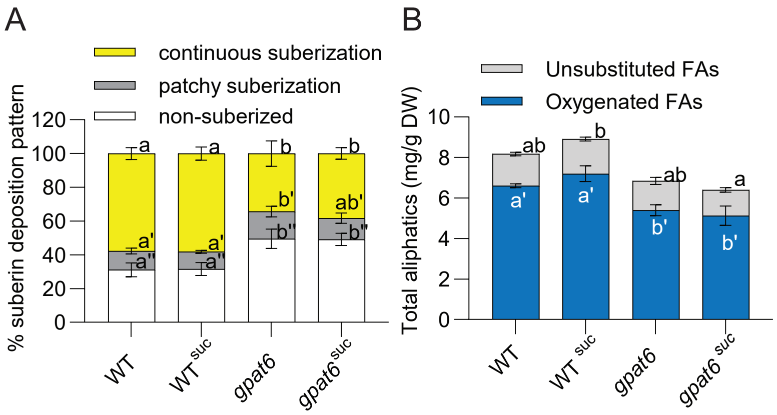


**Fig. S16. Suberin deposition in presence of low concentrations of sucrose**

(A-B) Since germination and seedling survival of *gpat4 gpat8* and *gpat4 gpat6 gpa8* mutants strongly depend on a low sucrose complement in the. MS plates, the suberization of different genotypes grown under standard conditions were compared to this in presence of 0.3 % sucrose (suc) in the medium. (A) The suberin pattern along the roots was evaluated by staining with Fluorol Yellow. n ≥ 6*;* values represent means +/- SD. Significant differences are shown with different letters by ANOVA, Tukey test, *p*-value < 0.05. (B) The suberin composition was evaluated. The total of the aliphatic suberin components group by substance classes are presented. Values represent the means ± SD, n =4. Significant differences to wild-type control shown with different letters by ANOVA, Tukey test *p*-value < 0.05.
