## Supplemental Movies description for "The *GPAT4*/*6*/*8* clade functions in Arabidopsis root suberization non-redundantly with the *GPAT5/7* clade required for suberin lamellae"

**Supplementary Movie 1**

Tomogram of the suberin lamellae in a Col-0 root, cut at 1mm from hypocotyl. Tilt series (-61°, +60°) through a Z volume of 118.64 nm. Scale bar: 200 nm.

**Supplementary Movie 2**

Reconstruction of the Tomogram from Supplementary Movie 1. Travelling in a series of 0.58 nm optical tomography slices through a Z volume of 118.64 nm showing the suberin lamellae in an endodermal cell of a Col-0 at 1 mm from the hypocotyl. Scale bar: 200 nm.

**Supplementary Movie 3**

Tomogram of the suberin lamellae in a Col-0 root treated with the phosphatase inhibitor PhosSTOP™ (0.5X), cut at 1 mm from hypocotyl. Tilt series (-61°, +60°) through a Z volume of 116.32 nm. Scale bar: 200 nm.

**Supplementary Movie 4**

Reconstruction of the Tomogram from Supplementary Movie 3. Travelling in a series of 0.58 nm optical tomography slices through a Z volume of 116.32 nm showing the suberin lamellae in an endodermal cell of a Col-0 treated with the phosphatase inhibitor PhosSTOP™ (0.5X) cut at 1 mm from the hypocotyl. Scale bar: 200 nm.

**Supplementary Movie 5**

Tomogram of the globular suberin in a *gpat5gpat7* mutant root, cut at 1 mm from hypocotyl. Tilt series (-56°, +54°) through a Z volume of 125.9 nm. Scale bar: 200 nm.

**Supplementary Movie 6**

Reconstruction of the Tomogram from Supplementary Movie 5. Travelling in a series of 0.58 nm optical tomography slices through a Z volume of 125.9 nm showing the absence of lamellated structures within the globular suberin in an endodermal cell of a *gpat5gpat7* mutant root at 1 mm from the hypocotyl. Scale bar: 200 nm.

**Supplementary Movie 7**

Tomogram of the suberin in a *gpat5gpat7* mutant root treated with the phosphatase inhibitor PhosSTOP™ (0.5X), cut at 1 mm from hypocotyl. Tilt series (-56°, +55°) through a Z volume of 153.84 nm. Scale bar: 200 nm.

**Supplementary Movie 8**

Reconstruction of the Tomogram from Supplementary Movie 7. Travelling in a series of 0.77 nm optical tomography slices through a Z volume of 153.84 nm showing the presence of lamellated structures within the suberin in an endodermal cell of a *gpat5gpat7* mutant root treated with the phosphatase inhibitor PhosSTOP™ (0.5X)at 1 mm from the hypocotyl. Scale bar: 200 nm.
