## Supplementary material for "The *GPAT4*/*6*/*8* clade functions in Arabidopsis root suberization non-redundantly with the *GPAT5/7* clade required for suberin lamellae": SMethod info and primer list

**Supplemental method information:**

**Sample fixation for transmission electron microscopy**

For chemical fixation, plants were fixed in 2.5% glutaraldehyde solution (Electron Microscopy Services) in 0.1 M phosphate buffer pH 7.4 for 1 h at room temperature and post-fixed in a fresh mixture of osmium tetroxide 1% (Electron Microscopy Services) with 1.5% potassium ferrocyanide (Sigma) in phosphate buffer for 1 h at room temperature. The samples were then washed twice in distilled water and dehydrated in acetone solution (Sigma) at graded concentrations (30%, 40 min; 50%, 40 min; 70%, 40 min; and 100%, 2 × 1 h). This was followed by infiltration in LR White resin (Electron Microscopy Science) at graded concentrations (LR White 33% in ethanol, 4 h; LR White 66% in ethanol, 4 h; LR White 100%, 2 × 8 h) and finally polymerized for 48 h at 60 °C in an oven under nitrogen atmosphere. For the higher-order mutants, 50-nm-thick sections were cut transversally 2 mm below the hypocotylroot junction, using a Leica Ultracut (Leica Mikrosysteme), picked up on a copper slot grid 2 × 1 mm (Electron Microscopy Science) coated with a polystyrene film (Sigma). For high-pressure freezing, plants were fixed in 2.5% glutaraldehyde solution (Electron Microscopy Services) in 0.1 M phosphate buffer pH 7.4 for 1 h at room temperature and post-fixed in a fresh mixture of osmium tetroxide 1% (Electron Microscopy Science) with 1.5% potassium ferrocyanide (Sigma) in phosphate buffer for 1 h at room temperature. The samples were then washed twice in distilled water before a high-pressure freezing step.

For the high-pressure freezing, 2-mm-long root pieces were cut below the hypocotyl junction region, and then placed in an aluminium planchet 3 mm in diameter with a 0.2 mm cavity (article 241, Wohlwend) filled with hexadecene (Merck) covered with a tap planchet (article 353, Wohlwend) and directly frozen under high pressure using a Leica Ice machine (Leica Microsysteme). The samples were then dehydrated and infiltrated with resin at low temperature using the Leica AFS2 freeze-substitution machine (Leica Mikrosysteme) with the following protocol: dehydration in 100% acetone (Sigma) at graded temperature (−90 °C, 10 h; −90 °C to −60 °C, 2 h; −60 °C, 8 h; −60 °C to −30 °C, 2 h; −30 °C, −3 h.) This was followed by infiltration in Spurr resin (Electron Microscopy Science) at graded concentration and temperature (30%, −30 °C to 0 °C, 10 h; 66%, 0 °C to 20 °C, 10 h; 100%, 20 °C, 2 × 10 h) and finally polymerized for 48 h at 60 °C in an oven. Fifty-nanometer sections were cut transversally to the root, using a Leica Ultracut (Leica Microsystem), picked up on a copper slot grid of 2 × 1 mm (Electron Microscopy Services) coated with a polystyrene film (Sigma).

For electron tomography, semi-thin sections of 200 nm thickness were cut transversally at 1 mm from the hypocotyl using a Leica Ultracut UC7 (Leica Mikrosysteme GmbH, Vienna, Austria) and then, picked up on a copper slot grid 2x1mm (EMS, Hatfield, PA, US) and coated with a polystyrene film (Sigma, St Louis, MO, US). Sections were post-stained with uranyl acetate (Sigma, St Louis, MO, US) 2% in $H_2O$ for 10 min and rinsed several times with $H_2O$.

Table 1

Primer used in this study

| Name | Sequence | Source |
|---|---|---|
| site-directed mutagenesis | | |
| GPAT4_mut_fw | GC CTC GGT AAA CGC ACC TCC ACA CAC GAT TTC | This study |
| GPAT4_mut_rev | CAA GAT CCG GTG AGT CAT CGC CGA ACT C | This study |
| GPAT6_mut_fw | GG CTC GGC AAA AGC AAG ACG ACA CAC GAC TTC | This study |
| GPAT6_mut_rev | CCA AAT CAG GTA AAT CAG ACG CTA GGC CAC | This study |
| GPAT8_mut_fw | GGC CTC GGT AAA CGA ACC TCT ACA CAT GAT TTC | This study |
| GPAT8_mut_rev | GAG ATC AGG TGA TTC GTT ACC AAA CTC TT | This study |
| Genotyping | | |
| GPAT4_mut_geno_fw | GTAGTGGTGACTGCGAATCC | This study |
| GPAT4_mut_geno_rev | CGG TTC TTG AGA CTC TCT ATA G | This study |
| GPAT6_mut_geno_fw | CTTCATCACGTTCGCGGGT | This study |
| GPAT6_mut_geno_rev | CGC ATT TCG TAC GTG GCA C | This study |
| GPAT8_mut_geno_fw | GATTGTGATGGTGGAGGCG | This study |
| GPAT8_mut_geno_rev | GCGGTTCTTTAAGCGTTCTTTTG | This study |
| GPAT4_LB | TCTCTTCCCATCGTCATCATC | Li *et al.* 2007 |
| GPAT4_RB | ACTGTTGTGGCTGATTTGGTC | Li *et al* . 2007 |
| GPAT5_LB | TTGGTTACTATATGCTCCTATTTTGG | Naseer *et al* . 2012 |
| GPAT5_RB | TTCGGACAAATGGTGAATTTC | Naseer *et al* . 2012 |
| GPAT6-1_LB | GTTGTAACGGGCGATACGTT | Li *et al* . 2012 |
| GPAT6-1_RB | CGTGACGTCGTTTTGAGAGA | Li *et al* . 2012 |
| GPAT6-2_RB | CACTTGAAAGGTTCCAACAAATC | Li *et al* . 2012 |
| GPAT7_LB | GTCGGAGCTAGAAGGAACACTAC | Naseer *et al* . 2012 |
| GPAT7_RB | CATAAACCGGTCTCGGGTTC | Naseer *et al* . 2012 |
| GPAT8_LB | GAACAGTACCTTGCAGAGAGACATGA | Li *et al.* 2007 |
| GPAT8_RB | TAATGAATTCGAACATGTGGCC | Li *et al* . 2007 |
| Promotor cloning | | |
| promGPAT1_fw | CAT AGG TAC CAC AAT GAC CGG GAG AAG | This study |
| promGPAT1_rev | CAT ACC CGG GAG CTA TGG CGT AGA GAG | This study |
| promGPAT2_fw | CATAGGTACCGTTCTCTGTTTTGGTCTTCTTG | This study |
| promGPAT2_rev | CATACCCGGGTTTGACCTCTCGTTTTCTAATAAC | This study |
| promGPAT3_fw | CATAGGTACCCCTGTTAGCTGGAGATGTTAGG | This study |
| promGPAT3_rev | CATACCCGGGGTTTGGATTTTGCAGAAAGC | This study |
| promGPAT4_fw | CATAGGTACCAACTTCATTGTTGCATCTTGG | This study |
| promGPAT4_rev | CATACCCGGGCTTTCTTGCGGCGAATACT | This study |
| promGPAT6_fw | CATAGGTACCGTCGTATTACAATGATGATCAAC | This study |
| promGPAT6_rev | CATACCCGGGAGATTTGGAAGGTGAGAATGG | This study |
| promGPAT7_fw | CATAGGTACCTGGGAAGATGTAGTCAAGCAC | This study |
| promGPAT7_rev | CATACCCGGGCACAACTTAACTTGTTTCTTTTTTTG | This study |
| promGPAT8_fw | CATAGGTACCTCAAGTTTTGGGATCTTCATG | This study |
| promGPAT8_rev | CATACCCGGGAACTATGAAATATCCACTAAGAGCG | This study |
| Real time PCR | | |
| SAND_qPCR_fw | AACTCTATGCAGCATTTGATCCACT | Hilfiker *et al* . 2014 |
| SAND_qPCR_rev | TGATTGCATATCTTTATCGCCATC | Hilfiker *et al* . 2014 |
| GPAT4_qPCR_fw | GTG TCA CTT ATA GTG TCT CTC GCC TC | This study |
| GPAT4_qPCR_rev | GCT CTG CAA ATAGAG CAC TGA ACC | This study |

| | | |
|---|---|---|
| GPAT5_qPCR_fw | TCGTTATGTGAGGAGCATATTCATG | This study |
| GPAT5_qPCR_rev | TTGTTGGTCACCGTGGTTGT | This study |
| GPAT6_qPCR_fw | CACCGAAGCAGAGAAGATAAAGG | This study |
| GPAT6_qPCR_rev | TAGGGTTCCGGTGAAGAAGG | This study |
| GPAT7_qPCR_fw | GGTCTAGGACGACATATTATCTCGG | This study |
| GPAT7_qPCR_rev | GCCGCTGGTTATGACCG | This study |
| GPAT8_qPCR_fw | GACTATGCGGTTCTTTAAGCGTTC | This study |
| GPAT8_qPCR_rev | GTTTGGTAACGAATCACCTGATCTCG | This study |
